# Decoding the Epigenetics and Chromatin Loop Dynamics of Androgen Receptor-Mediated Transcription

**DOI:** 10.1101/2023.12.22.573134

**Authors:** Umut Berkay Altıntaş, Ji-Heui Seo, Claudia Giambartolomei, Dogancan Ozturan, Brad J. Fortunato, Geoffrey M. Nelson, Seth Raphael Goldman, Karen Adelman, Faraz Hach, Matthew L. Freedman, Nathan A. Lack

## Abstract

Androgen receptor (AR)-mediated transcription plays a critical role in normal prostate development and prostate cancer growth. AR drives gene expression by binding to thousands of cis-regulatory elements (CRE) that loop to hundreds of target promoters. With multiple CREs interacting with a single promoter, it remains unclear how individual AR bound CREs contribute to gene expression. To characterize the involvement of these CREs, we investigated the AR-driven epigenetic and chromosomal chromatin looping changes. We collected a kinetic multiomic dataset comprised of steady-state mRNA, chromatin accessibility, transcription factor binding, histone modifications, chromatin looping, and nascent RNA. Using an integrated regulatory network, we found that AR binding induces sequential changes in the epigenetic features at CREs, independent of gene expression. Further, we showed that binding of AR does not result in a substantial rewiring of chromatin loops, but instead increases the contact frequency of pre-existing loops to target promoters. Our results show that gene expression strongly correlates to the changes in contact frequency. We then proposed and experimentally validated an unbalanced multi-enhancer model where the impact on gene expression of AR-bound enhancers is heterogeneous, and is proportional to their contact frequency with target gene promoters. Overall, these findings provide new insight into AR-mediated gene expression upon acute androgen simulation and develop a mechanistic framework to investigate nuclear receptor mediated perturbations.

## INTRODUCTION

The androgen receptor (AR) is a ligand-dependent transcription factor that plays a critical role in regulating gene expression in the prostate^1^. In its inactive form, the AR resides in the cytoplasm where it is stabilized by heat-shock chaperone proteins^2^. After binding androgens, such as testosterone or dihydrotestosterone (DHT), the AR undergoes an allosteric modification and translocates into the nucleus^2–4^. Once there, the AR binds to specific cis-regulatory elements (CREs) on DNA through an interplay of chromatin accessibility, pioneer factors such as FOXA1, and sequence motifs^5–9^. The majority of these AR-bound CREs are proposed to function as enhancers as they are located distal from gene promoters^10^ and are brought into close physical proximity by chromatin loops^11,12^. The enhancer activity at these CREs is associated with various epigenetic features, including chromatin accessibility, transcription factor binding, and post-translational histone modifications, such as H3K27ac and H3K4me3^13–19^. CREs are proposed to impact transcription through chromatin contacts with the target promoter that cause the recruitment of co-regulatory proteins and transcriptional machinery^20–23^. This typically involves multiple AR-bound enhancers and CREs that interact with the target promoter^24–30^. The contribution of individual CREs has been controversial with some studies suggesting that CREs work additively to increase gene expression^31–33^, while others propose that many enhancers are redundant and only contribute at specific developmental stages^34–37^. The complex kinetic interplay of epigenetic modifications, co-regulatory proteins and chromatin loops across multiple CREs following transcription factor binding is poorly understood.

To explore how AR binding impacts epigenetic modifications and chromatin looping at regulatory elements in response to an acute perturbation, we generated an extensive multiomics experimental dataset following androgen stimulation that was integrated into a graph-based framework. We demonstrated that AR binding sequentially induces an increase in FOXA1 and H3K27ac signals, that was followed by an increase in chromatin accessibility. We further show that AR does not induce new chromatin loops, but instead increases the contact frequency between gene promoters and selective AR-bound enhancers. From these results, we generated and validated a multi-enhancer model, where a small subset of pre-established dominant CREs with increased chromatin contact frequency exhibits an elevated dynamic response to androgen stimulation, which significantly contribute to gene expression. These results provide a foundation for understanding how enhancers respond to an acute perturbation.

## METHODS

### Experimental

#### LNCaP Cell culture and DHT treatment

LNCaP cells (#CRL-1740, ATCC) were grown in phenol red-free RPMI (#11835030, GIBCO) with 10% charcoal-stripped FBS (#100-119, GemBio) for 3 days and then were stimulated with 10nM dihydrotestosterone (DHT) (A8380, Sigma) for 0.5, 4, 16 or 72 hours. For the vehicle samples, the cells were treated with 100% EtOH. Subsequently, cells were collected for further experiments (ChIP-seq, RNA-seq, ATAC-seq, HiChIP, or Start-seq) accordingly^38^. LNCaP cells were authenticated by sequencing and comparing short tandem repeats to parental LNCaP cells in the ATCC database. Prior to experiments, cells were tested for mycoplasma contamination with LookOut Mycoplasma PCR Detection Kit (Sigma-Aldrich #D9307).

#### Chromatin Immunoprecipitation Assays with Sequencing (ChIP-seq)

ChIP-seq in LNCaP was performed as previously described^38^. Briefly, 10 million cells were fixed with 1% formaldehyde at room temperature for 10 min, quenched, and collected in lysis buffer (1% NP-40, 0.5% sodium deoxycholate, 0.1% SDS and protease inhibitor [#11873580001, Roche] in PBS). Chromatin was sonicated with a Covaris E220 sonicator (140PIP, 5% duty cycle, 200 cycle burst). The sample was then incubated with antibodies (AR: Abcam ab133273, FOXA1: Abcam ab23738, H3K27ac: Diagenode 3C15410196; H3K4me3: Diagenode 3C15410003) coupled with Dynabeads protein A and protein G beads (Life Technologies) at 4 °C. Incubated chromatin was washed with RIPA wash buffer (100 mM Tris pH 7.5, 500 mM LiCl, 1% NP-40, 1% sodium deoxycholate) for 10 min six times and rinsed with TE buffer (pH 8.0) once. DNA was purified using a MinElute column followed by incubation in the de-crosslinking buffer (1% SDS, 0.1 M NaHCO3 with Proteinase K and RNase A) at 65 °C. Eluted DNA was prepared as sequencing libraries with the ThruPLEX-FD Prep Kit (Takara bio, # R400675). Libraries were sequenced with 150-BP PE on an Illumina HiSeq 2500 Sequencing platform at Novogene.

#### RNA-Seq

LNCaP cells (5×10^5) were harvested for RNA-seq^39^. Total RNA was collected from the cells using an RNeasy kit (Qiagen, #74104,) with DNase I treatment (Qiagen, #79254). The library preparations, quality control, and sequencing on HiSeq 2500 Sequencing platforms (150-BP PE) were performed by Novogene.

#### Assay for transposase-accessible chromatin with sequencing (ATAC-seq)

LNCaP cells were isolated and subjected to modified ATAC–seq as previously described^38^. Briefly, 50,000 nuclei were pelleted and resuspended in ice-cold 50μl of lysis buffer (10 mM Tris-HCl, pH 7.5, 10mM NaCl, 3mM MgCl2, 0.1% NP-40, 0.1% Tween20, and 0.01% Digitonin). The subsequent centrifugation was performed at 500 g for 10 min at 4 °C. The nuclei pellets were resuspended in 50μl of transposition buffer (25μl of 2× TD buffer, 22.5μl of distilled water, 2.5 μl of Illumina Tn5 transposase) and incubated at 37 °C for 30 min with shaking at 1000 rpm for fragmentation. Transposed DNA was purified with the MinElute PCR Purification kit (Qiagen). DNA was purified using Qiagen MinElute (#28004), and the library was amplified up to the cycle number determined by 1/3rd maximal qPCR fluorescence. ATAC-seq libraries were sequenced with 150-BP PE high-throughput sequencing on an Illumina HiSeq 2500 Sequencing platform (Novogene).

#### HiC combined with capture ChIP-seq (HiChIP)

HiChIP was performed as previously described^39^. Trypsinized LNCaP cells (10 million) were fixed with 1% formaldehyde at room temperature for 10 min and quenched. The sample was lysed in HiChIP lysis buffer and digested with MboI (NEB) for 4 h. After 1 h of biotin incorporation with biotin dATP, the sample was ligated with T4 DNA ligase for 4 h with rotation. Chromatin was sonicated using Covaris E220 (conditions: 140 PIP, 5% DF, 200 CB) to 300-800 bp in ChIP lysis buffer (1% NP-40, 0.5% sodium deoxycholate, 0.1% SDS and protease inhibitor in PBS) and centrifuged at 13,000 rpm. for 10 minutes at 4 °C. Preclearing 30μl of Dynabeads protein A/G for 1 h at 4 °C was followed by incubation with antibodies (H3K27ac, Diagenode, 3C15410196; H3K4me3, Diagenode 3C15410003). Reverse-crossed IP sample were pulled down with streptavidin C1 beads (Life Technologies), treated with Transposase (Illumina) and amplified with reasonable cycle numbers based on the qPCR with a 5-cycle pre-amplified library. The library was sequenced with 150-BP PE reads on the Illumina HiSeq 2500 Sequencing platform (Novogene).

#### Small Capped Nascent RNA Sequencing (Start-seq)

For Start-seq, LNCaP cells were grown and collected as described as above. Cell pellets were washed with ice-cold 1x PBS. 1 million cells were treat with 1.5mL of Nun Buffer (0.3M NaCl, 1M Urea, 1% NP-40, 20mM HEPES pH 7.5, 7.5mM MgCl2, 0.2M EDTAm, protease inhibitors and 20U/mL SUPERase-IN) for 30 min on ice with frequent vortexing. Chromatin was pelleted by centrifugation at 12500 g for 30 min for 4 °C. After 3 times-washing with 1mL ice-cold chromatin washing buffer (50mM Tris-HCL pH7.5 and 40U/mL SUPERase-In) and additional 0.5mL of Nun buffer. After centrifugation for 5 min at 500 g, 0.5m TRIzol was added to the remaining pellet. Libraries were prepared according to the TruSeq Small RNA Kit(Illumina). To normalize samples, 15 synthetic capped RNAs were spiked into the Trizolpreparation at a specific quantity per 10^7 cells, as in^40^. The library was sequenced with 150-BP PE reads on the Illumina HiSeq 2500 Sequencing platform (Novogene).

#### CRISPRi

Guide RNAs (gRNAs) were designed for each enhancer and promoter region using CRISPR-SURF^41^. Cas-OFFinder was used to eliminate off-target gRNAs^42^. LNCaP cells stably expressing dCas9-KRAB (Addgene #89567) were seeded in a 6-well plate at a density of 200K cells per well. For transfection, a total of 500-1500ng DNA was used, divided according to the number of available gRNAs. Transfection was performed using Mirus TransIT-X2. After transfection, the media was replaced and 2ng/µl of puromycin was added for selection. Following 72 h of selection, the medium was changed to charcoal-stripped serum for androgen deprivation. After 48 h, the cells were treated with 10nM DHT for 4 h and then trypsinized. RNA extraction and cDNA preparation was performed using the LunaScript® RT SuperMix Kit. The androgen-induced expression was quantified using qRT-PCR and each experiment was conducted in triplicate. The sequences of gRNAs and qRT-PCR primers can be found in **Supp Table 1**.

### Bioinformatics Analyses

#### RNA-seq analysis

Reads were aligned to the hg19 human genome with STAR (v2.7.9)^43^ with quant mode (-- quantMode TranscriptomeSAM). Next, “toTranscriptome” bam files were fed into Salmon (v0.14.1)^44^ to quantify TPM values for (Gencode v19)^45^ transcripts. To generate the signal track files from RNA-seq, VIPER^46^ is used. Briefly, STAR as the default aligner, converts BAM files into BigWig files using Bedtools (v2.30.0)^47^.

#### ChIP-seq and ATAC-seq analyses

ChIP-seq and ATAC-seq were processed through the ChiLin pipeline^48^. Briefly, Illumina Casava1.7 software used for base calling and raw sequence quality and GC content were checked using FastQC (v0.10.1). The Burrows–Wheeler Aligner (BWA^49^, v0.7.10) was used to align the reads to the human genome hg19. Then, MACS2^50^ (v2.1.0.20140616) was used for peak calling with an FDR q-value threshold of 0.01. Bed files and Bigwig files were generated using bedGraphToBigWig^51^ and the union of narrow and broad peaks from ChIP-seq were used as anchors to call loops. The following quality metrics were assessed for each sample: (i) percentage of uniquely mapped reads, (ii) PCR bottleneck coefficient to identify potential over-amplification by PCR, (iii) FRiP (fraction of non-mitochondrial reads in peak regions), (iv) peak number, (v) number of peaks with 10-fold and 20-fold enrichment over the background, (vi) fragment size, (vii) the percentage of the merged peaks with promoter, enhancer, intron, or intergenic, and (viii) peak overlap with DNaseI hypersensitivity sites. For datasets with replicates, the replicate consistency was checked by two metrics: 1. Pearson correlation of reads across the genome using UCSC software wigCorrelate after normalizing signal to reads per million and 2. percentage of overlapping peaks in the replicates.

#### HiChIP Loop calling

After trimming adaptors from the HiChIP datasets using Trim Galore (v0.5.0) (https://github.com/FelixKrueger/TrimGalore), we used HiC-Pro (v3.1.0)^52^ as previously described in Giambartolomei and Seo *et al.*^39^. This aligned the reads to the hg19 human genome, assigned reads to MboI restriction fragments, and removed duplicate reads. We used the following options: <MIN_MAPQ = 20, BOWTIE2_GLOBAL_OPTIONS = –very-sensitive– end-to-end–reorder, BOWTIE2_LOCAL_OPTIONS = –very-sensitive–end-to-end–reorder, GENOME_FRAGMENT = MboI_resfrag_hg19.bed, LIGATION_SITE = GATCGATC, LIGATION_SITE = “GATCGATC,” BIN_SIZE = “5000.”> All other default settings were used. To build the contact maps, the HiC-Pro pipeline selects only uniquely mapped valid read pairs involving two different restriction fragments. We applied FitHiChIP (v10.0)^53^ for bias-corrected peak and DNA loop calls. FitHiChIP models the genomic distance effect with a spline fit, normalizes for coverage differences with regression, and computes statistical significance estimates for each pair of loci. We used the FitHiChIP loop significance model to determine whether interactions are significantly stronger than the random background interaction frequency. As anchors to call loops in the HiChIP analyses, we used 842367 regions for H3K27ac and 136939 regions for H3K4me3, which resulted from merging ChIP-seq narrow and broad peaks comprised 2 replicates for each broad and narrow peak and each of the five-time points (0m, 30m, 4h, 16h, 72h) for H3K27ac, and 1 replicate for each broad and narrow peak and each of the same time points for H3K4me3. We used a 5 kb resolution and considered only interactions between 5 kb and 3 Mb. We used the peak to all for the foreground, meaning at least one anchor needed to be in the peak rather than both. The corresponding FitHiChIP options specification is <IntType=3> For the global background estimation of expected counts (and contact probabilities for each genomic distance), FitHiChIP can use either peak-to-peak (stringent) or peak-to-all (loose) loops for learning the background and spline fitting. We specified the suggested option to merge interactions close to each other to represent a single interaction when their originating bins are closer. The corresponding FitHiChIP options specifications are <UseP2PBackgrnd=0> and <MergeInt=1> (FitHiChIP (L + M)). We used the default FitHiChIP q value < 0.01 to identify significant loops. We used hicpro2higlass (v3.1.0) to convert allValidPairs to .cool files after having specified the hg19 chromosome sizes, using the following command: <hicpro2higlass.sh -i sample.allValidPairs -r 5000 -c chrsizes -n> The reads from HiChIP data were merged from every time point for H3K27ac and H3K4me3 separately, and the reference loop sets were called with the same parameters above (resulting loop number, n=296,326; n=278,491 respectively). Next, the contact frequency values of each time point at loops are captured from the .cool files using the Python package, Cooler (v0.9.3). The count table was then TMM normalized among time points to reduce the between-sample variation.

#### Annotation of Cis-Regulatory Elements / Defining Background Genes

CREs were defined as +/-2.5 kb around the summit of the accessible region from ATAC-seq peaks at any given time point. Promoters were defined with a multi-step process. First, we identified the highest expression transcript (Gencode v19)^45^ isoforms using Salmon (v0.14.1)^44^, see RNA-seq analysis. Next, the start locations -according to strand information-of the highest expressing transcripts were collected and the summit was extended +/2.5 kb. If they overlap with a defined CRE, we assigned the overlapping CRE as the active promoters. From these annotated active promoters, the genes of HALLMARKS_ANDROGEN_RESPONSE from the Molecular Signature Database (MSigDB)^54^ were defined as AR-regulated genes. The transcriptome was divided into four quartiles and AR-regulated genes were compared with similarly expressing genes (**Supp Fig 1A, B**). Overlaping CREs to an AR ChIP-seq peaks at any time point were defined as potential AR-bound enhancers. The median number AR-bound enhancers (E+AR; ∼2) of AR-regulated genes were less than AR-free enhancers (E-AR; ∼14) (**Supp Fig 1E**). Similarly, a CRE was considered FOXA1-bound, if it intersects with a FOXA1 peak at any time point (**Supp Fig 1C**). To identify down-regulated genes, we calculated the log2foldChange induction between 16h and 0m, considering those with a value below -1 as downregulated genes (**Supp Fig 3A, B**).

#### Constructing the Graph Network

When we called significant loops from each time point separately, there were few called loops overlapping between time points potentially due to both the loop calling methodologies and experimental noise present in HiChIP data^55^. As we could observe the matching loops in the contact matrices of all time points (**Supp Fig 5**), we instead generated a reference loop set, normalized the count matrix, and compared contact frequencies similar to published work^20^. A custom R (v4.1.1) script (https://github.com/lacklab/AR_transcription) was used to extract interaction pairs of annotated CRE regions (BED) according to the reference loop set (BEDPE; both H3K27ac and H3K4me3) using ‘GenomicInteractions’ (v1.34)^56^. The graph structure is built in custom Python (v3.9.7) script (link) using the NetworkX (v3.1) package^57^. Briefly, any pair of CREs are included as nodes in the network with TMM normalized HiChIP contact frequency as weighted edges, for both H3K27ac and H3K4me3 at every time point.

#### Epigenetic changes of CREs and kinetic changes of enhancers of AR-regulated genes

The average signal (AR, FOXA1, H3K27ac, H3K4me3, ATAC) at the promoters and CREs were collected using a custom python package (https://github.com/breambio/bluegill). To reduce the batch effect across time, TMM normalization was applied individually for each epigenetic factor^58^. To visualize the temporal change upon androgen stimulation, line plots were drawn (**Fig 2**). For the selected six AR-regulated genes (**Fig 3F**), both the average signal (AR, FOXA1, H3K27ac, H3K4me3, ATAC) over the first-degree interacting enhancers of each gene promoter and contact frequency (CF) between every gene promoter and corresponding enhancer were collected for every time point. The standard deviation (SD) of each feature along the time domain was calculated and min-max normalized.

**Figure 1:**
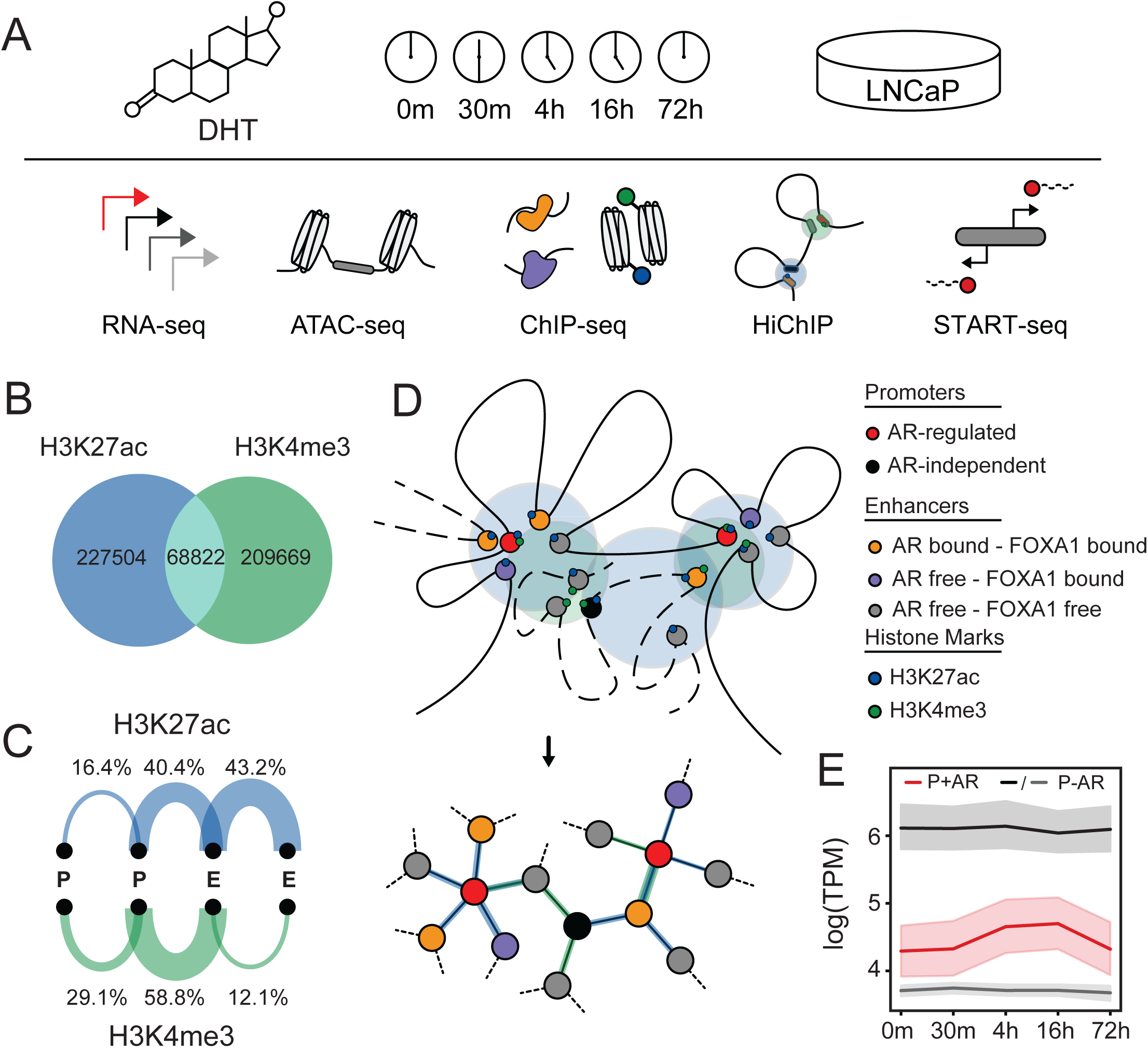
Design of temporal multi-omics dataset and construction of a bioinformatic framework. **(A)** Schematic representation of the experimental design. LNCaP cells were treated with 10nM DHT and samples were collected at 5 different time points (0m, 30m, 4h, 16h, 72h) for RNA-seq, ATAC-seq, ChIP-seq (AR, FOXA1, H3K27ac, H3K4me3), HiChIP (H3K27ac, H3K4me3) and Start-seq. **(B)** Venn diagram representing significantly called chromatin loops from merged H3K27ac and H3K4me3 HiChIP datasets. **(C)** Arc plots representing percentages of promoter-promoter (P-P), enhancer/CRE-promoter (E-P), and enhancer/CRE-enhancer/CRE (E-E) loops for H3K27ac and H3K4me3 HiChIP. **(D)** Graphical representation of regulatory network. **(E)** Gene expression profile of androgen response hallmark genes (P+AR; red), highly expressed (first quartile; P-AR; black) and mid-high (second quartile; P-AR; grey) expressed genes at all time points.

**Figure 2:**
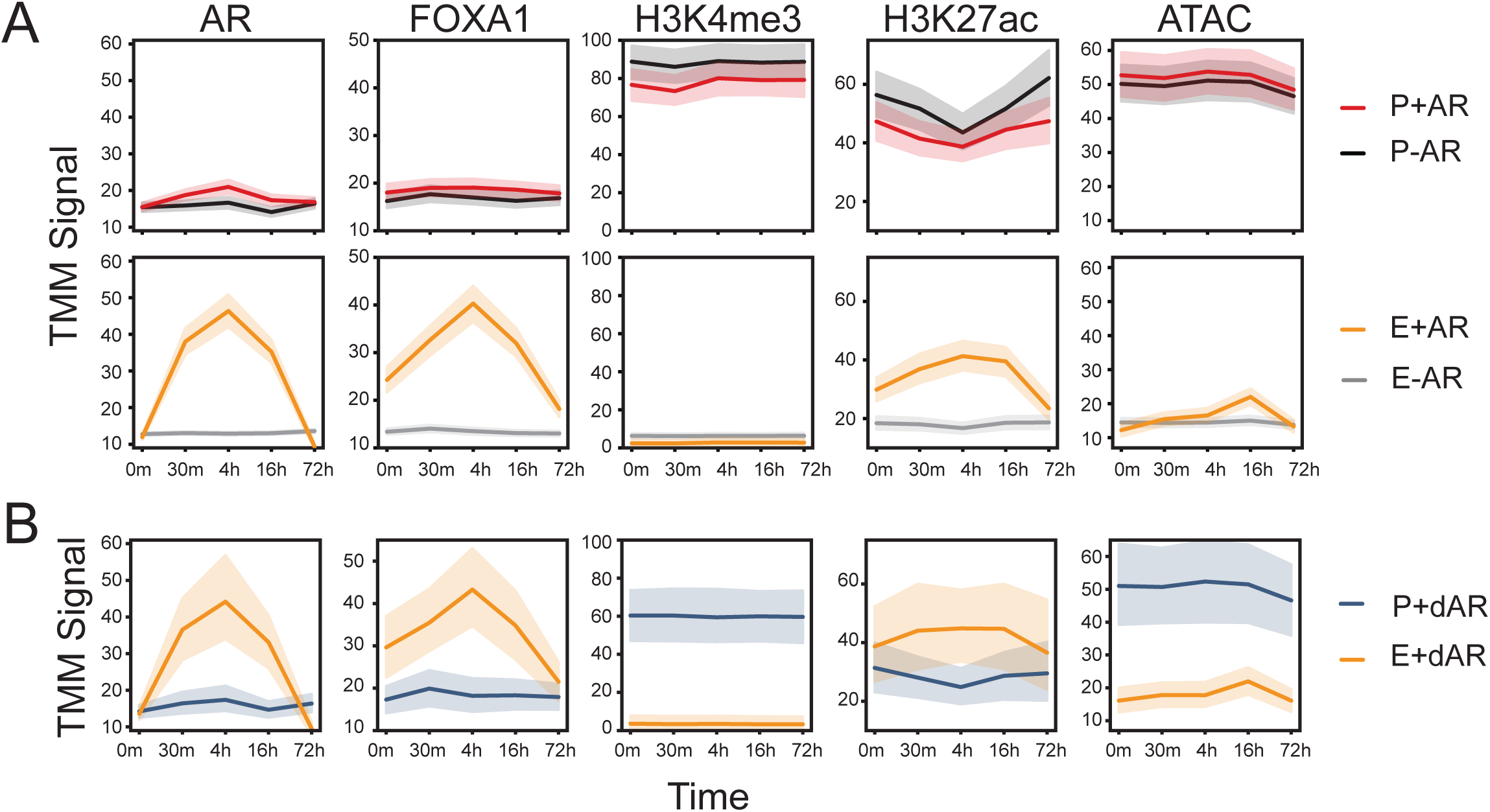
Activation of the androgen receptor leads to a delayed increase in histone modifications and chromatin accessibility. **(A)** TMM normalized ChIP-seq or ATAC-seq signal were compared across all time points at different regulatory elements including: Promoters of AR up-regulated genes (P+AR: red), promoters of AR-independent genes (P-AR: black), AR-bound enhancers (E+AR: yellow) of AR-regulated genes, AR-free enhancers (E-AR: grey) of AR-independent genes. **(B)** A similar analysis was done for the promoters of AR down-regulated genes (P+dAR: blue), and their AR-bound enhancers (E+dAR: yellow).

**Figure 3:**
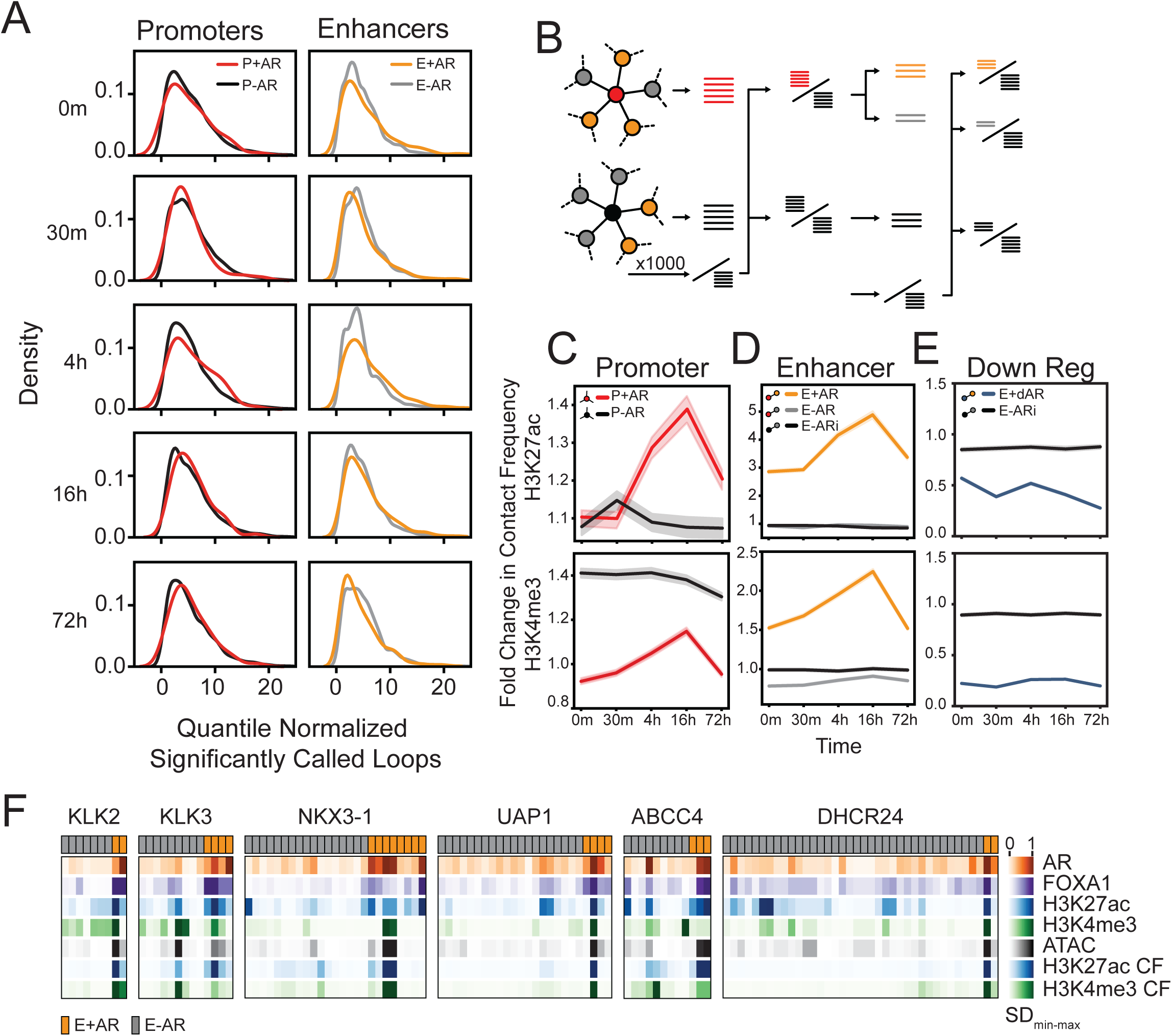
AR-bound enhancers increase contact frequency to AR-regulated gene promoters to upregulate gene expression. **(A)** Kernel density estimation of the number of significantly called loop anchoring promoters (left; AR-regulated promoters are in red; AR-independent promoter background are in black) or enhancers (right; AR-bound enhancers are in yellow; AR-free CRE background are in grey) at each time point (H3K27ac HiChIP). Each row represents the significantly called loops at each time point separately, and not reference loops. **(B)** Schematic representation of fold change in contact frequency calculation for a given gene set (left), from its promoter’s viewpoint (middle), from its enhancers’ viewpoint (right). The query loop sets are compared to the same reference loop set. Promoter view: The AR-regulated gene promoter loops (red sticks) and a random set of AR-independent gene promoter loops (black sticks) are compared to the reference loops which are selected from AR-independent promoter loops (denominator black sticks). Enhancer view: The loops to AR-bound enhancers (yellow sticks) and AR-free enhancers (grey sticks) and loops of AR-free enhancers (black sticks) that interacting with AR-independent gene promoters are compared to the reference loops which are selected from AR-independent promoters that interact with AR-free enhancers (denominator black sticks). **(C)** Fold change in chromatin loop contact frequency of AR-regulated gene promoters (P+AR; red) and highly expressed AR-independent gene promoters (P-AR; black). **(D)** Fold change in contact frequency of AR-bound enhancers that loop to AR-regulated genes (E+AR; yellow), AR-free enhancers looping to AR-regulated genes (E-AR; grey), and AR-free enhancers looping to highly expressed AR-independent genes (E-ARi; black) **(E)** Fold change in contact frequency in AR-bound enhancers looping to AR-down-regulated genes (E+dAR; blue), and AR-free enhancers looping to highly expressed genes (E-ARi; black) **(F)** Min-max normalized standard deviation (SD) following androgen treatment for AR, FOXA1, H3K27ac and H3K4me3 ChIP-seq, ATAC-seq and H3K27ac and H3K4me3 HiChIP contact frequency (CF) are depicted for AR-bound (E+AR; orange) and AR-free (E-AR; grey) CREs of six selected AR-regulated genes (*KLK2, KLK3, NKX3-1, UAP1, ABBCC4, DHCR24*).

#### Calculation of chromatin contact frequency change

Promoter-centric analysis was done with the query loop sets, AR-regulated genes’ promoter loops (P+AR), and random highly expressed genes’ promoter loops (P-AR; n=100 ; seed=7). This was compared to reference loop sets (n=100) that was randomized (1000 iterations; seed=7) from highly expressed genes’ promoter loops (**Fig 3B**). Fold change was calculated for each interation between the average contact frequency of loops in the query and reference. The same approach was used to calculate the contact frequency fold change from the enhancer viewpoint. The loop sets were selected according to the E-P pairs. While selecting the query and reference loops, the number of randomly selected promoters was fixed (n=100). The loops between AR-regulated gene promoters to AR-bound (E+AR) and AR-free enhancers (E-AR), and between AR-independent gene promoters to AR-free enhancers (E-AR) were compared to randomized loops from an AR-free enhancer to highly expressed gene promoters (1000 iterations; seed=7).

#### Start-seq analysis

Adaptor sequences and low-quality 3′ ends were removed from paired-end reads of all samples using cutadapt (v1.2.1). Reads shorter than 20 nucleotides were discarded (-m 20 -q 10), and a single nucleotide was trimmed from the 3′ end of all remaining reads to enable successful alignment with bowtie (v1.1.1). The first in pair flagged reads were filtered to generate signal (bigwig) files for each strand using bedtools genomecov (-bg -5 -strand +/-, respectively).

The transcripts (Gencode v19) were extracted GTF file. The maximum signal within a range of +/-500 bp of TSS was gathered from the forward Start-seq track for plus-stranded transcripts, and the reverse for minus-stranded transcripts. Next, the log2foldchange (LFC), compared to 0m, was calculated for every time point (30m, 4h, 16h, 72h). If a transcript was found to be LFC > 1 at any time point, it was considered as differential up regulation. Next, the nascent expression levels at all time points for those transcripts (the union of differential up-regulation) were z-normalized to capture the highest expression time point, which is assigned to time-based expression groups (**Fig 4A**). Similar to enhancer viewpoint analysis, the first-degree AR-bound enhancer contact frequency to the gene promoters was compared to randomly selected contact frequencies (1000 iterations) between the highly expressed gene promoters (n=100) and their first-degree AR-free enhancers.

**Figure 4:**
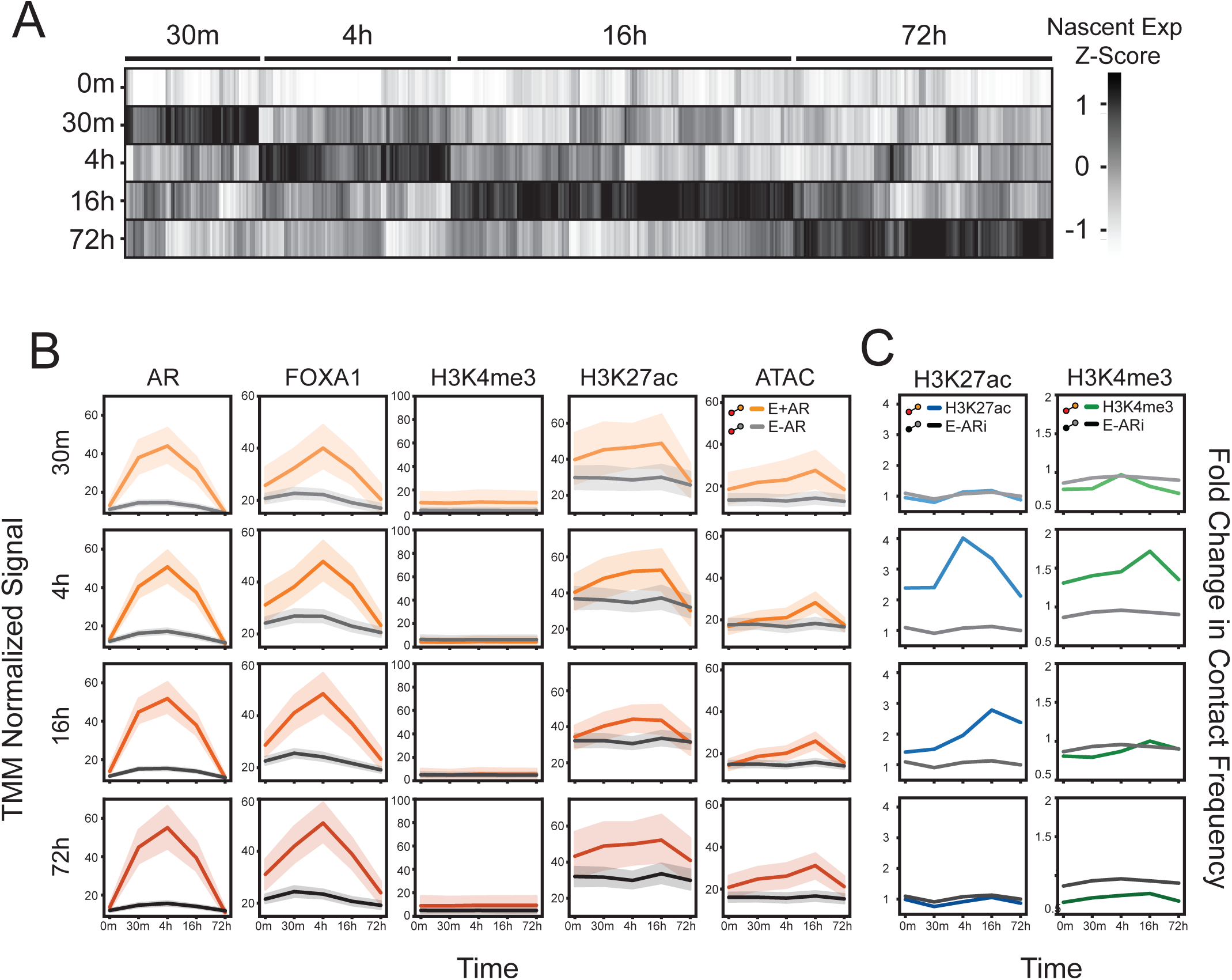
Temporal changes on epigenetics nascent RNA. **(A)** Clustering of nascent capped mRNA (Start-seq) according to time-dependent maximum expression. The union of upregulated transcripts’ nascent expression level across all time points were z-score normalized. **(B)** TMM normalized ChIP-seq or ATAC-seq of the following regulatory elements: AR-bound enhancers (E+AR: yellow) and AR-free enhancers (E-AR: grey). Each row represents the enhancers that loop to those genes that are maximally expressed at that time point (see A). **(C)** Fold change in contact frequency of AR-bound enhancers looping to maximally expressed genes for H3K27ac (E+AR; blue) and H3K4me3 (E+AR; green) HiChIP. These CREs were compared to AR-free enhancers of highly expressed genes (E-ARi; black) in both HiChIP datasets.

#### Contact frequency, expression correlation

To explore the contact frequency vs. expression, we performed three first-degree summarization models (average, maximum, and sum). Briefly, we identified all first-degree contacts of every gene promoter, then we summarized the contact frequencies with the corresponding function of the models for every gene. To reduce the signal noise, we first sorted genes according to expression, binned them into equal-sized sets (k=25), plotted bins as a scatter plot with average log expression on the x-axis, and averaged summarized contact frequency on the y-axis (**Fig 5B, Supp Fig 7**).

**Figure 5:**
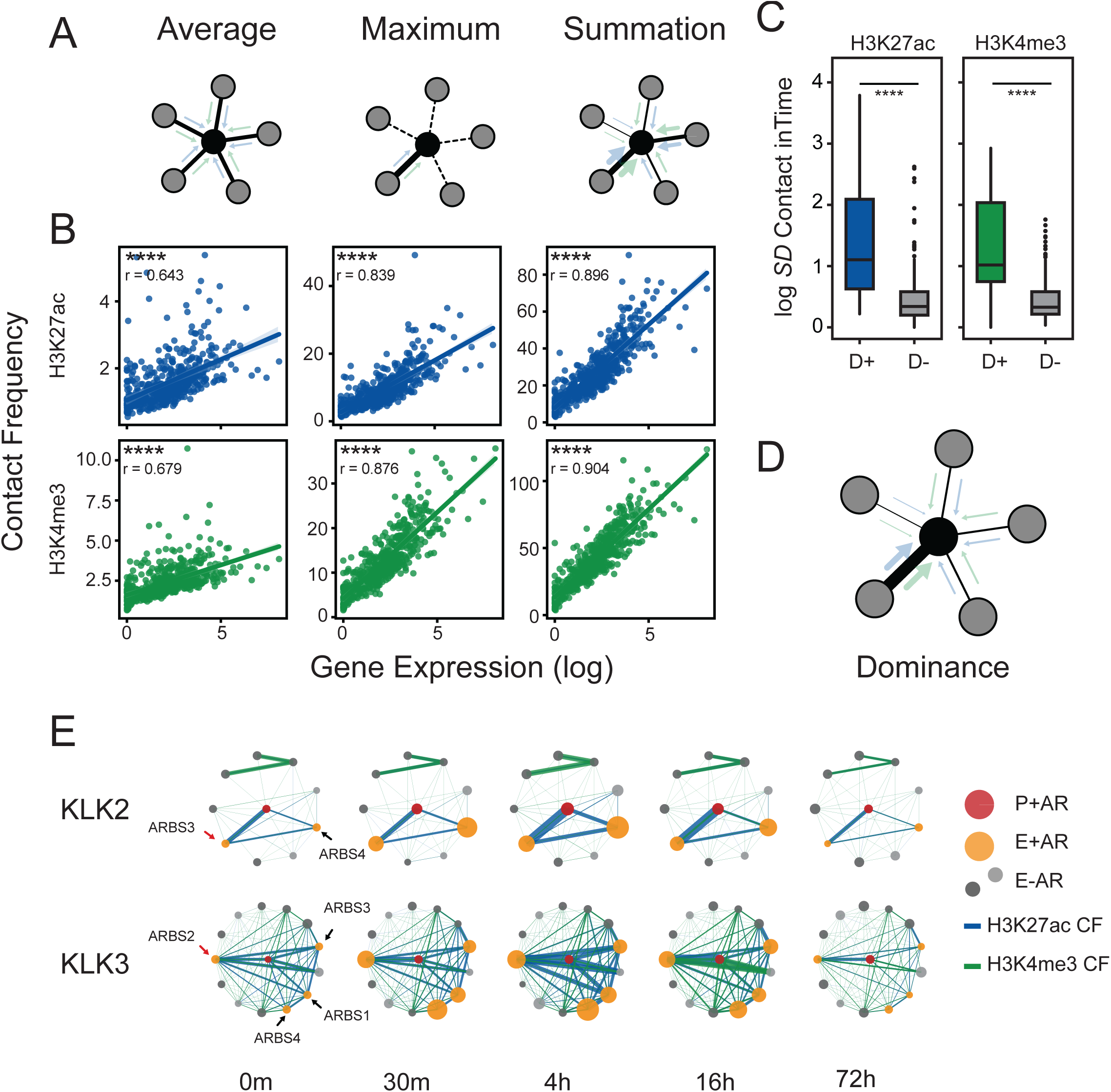
Multi-enhancer contact is strongly influenced by a dominant loop. **(A)** Schematic representation of tested multi-enhancer models. Average model (left) represents equal impact of CREs (surrounding nodes; grey) on gene promoter (center node; black). Maximum model (middle) represents only one neighbor (unbroken line) impact on gene promoter (center node; black). Summation model (right) accept the variability (different width of blue/green arrows) and represents all CREs impact additively with their contact frequency to gene promoter. **(B)** Scatter plot of binned (k=25) log expression (x-axis) vs. contact frequency (y-axis) according to the proposed multi-enhancer models function (16h HiChIP). Correlation was calculated by linear regression. **(C)** Standard deviation of H3K27ac and H3K4me3 HiChIP contact frequency change over time for dominant AR-bound CREs (D+) and non-dominant AR-bound CREs (D-) that interact with promoters of AR-regulated genes. **(D)** Schematic representation of hypothesized multi-enhancer dominance model. The width of arrows (blue/green) represent the contact frequency. **(E)** Circle plot representation of first-degree regulatory interactions of *KLK2* (top) and *KLK3* (bottom) genes. Promoters are shown in the centre (red) and the first-degree interactions are either AR-bound (yellow) or AR-free (greys) CREs. The size of the nodes represents the AR ChIP-seq signal, and the width of the edges represents contact frequency (H3K27ac CF: blue, H3K4me3 CF: green). For all data ns p>0.05, * p<0.05, ** p<0.01, *** p<0.001, **** p<0.0001.

#### Random Forest Regressor

The binned (k=25) contact frequency and expression data (see above) were used to train a random forest regressor from the Sklearn (v1.3.1)^59^ with “random_state=0”. The permutation importance of each feature is also calculated within the Sklearn^60^.

#### Gini Index Analysis

The chromatin contact frequency between gene promoter and first-degree interacting elements was identified and the Gini index was calculated for a 16h time-point (**Fig 5C**). To calculate the Gini index a custom python function was utilized (https://github.com/lacklab/AR_transcription). The mean absolute difference is the average absolute difference between all pairs of items in the population. The relative mean absolute difference is obtained by dividing the mean absolute difference by the population’s average (x) to account for differences in scale (e.q 1), where x_i_ is contact frequency of a loop *i*, and there are *n* loops of a promoter.

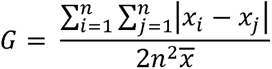

#### Circle Plots

Circle plots were drawn with NetworkX and Matplotlib libraries^57,61^ (https://github.com/lacklab/AR_transcription)

#### Statistical analysis

The probability of finding *k* loops from each CREs was determined by hypergeometric distribution (**Supp Fig 4**). When comparing the target set with the background set, a non-parametric Mann-Whitney-U test was applied. Correlation coefficients and their p-values were calculated with Spearman’s test. All statistical tests were performed using the SciPy (v1.11.1) Python package^62^.

## RESULTS

### Generation of androgen-stimulation kinetic dataset and construction of regulatory networks

To characterize the temporal impact of AR binding on epigenetic features and chromatin looping, we generated an extensive kinetic multi-omics experimental dataset following androgen stimulation. We treated LNCaP cells with dihydrotestosterone and collected cells at 5 different time points (0m, 30m, 4h, 16h, 72h). At each time point, multiple features were characterized including gene expression (RNA-seq), chromatin accessibility (ATAC-seq), transcription factor binding and post-translational histone modification (AR, FOXA1, H3K27ac and H3K4me3 ChIP-seq), chromatin looping (HiChIP) and capped nascent RNA (Start-seq) (**Fig 1A**). From these datasets, CREs (n=78,522) were defined from accessible sites (ATAC-seq)^15^. Based on known gene annotations and AR ChIP-seq, CREs were annotated as either promoters (n=13,575), AR-free CREs (n=58,265), or AR-bound CREs (n=6,682). The interaction between these CREs was defined from consensus H3K27ac (enhancer-centric; n=296,326) and H3K4me3 (promoter-centric; n=278,491; see methods) HiChIP loop sets. Supporting that these histone marks are associated with different functional CREs, only 23% of loops were found in both H3K27ac and H3K4me3 HiChIP (**Fig 1B**). As expected, H3K27ac loops were predominantly between enhancer-enhancer (E-E) pairs, accounting for 43.2% of the loops, followed by enhancer-promoter (E-P) pairs, which constituted around 40.4% (**Fig 1C**). In contrast, H3K4me3 loops were primarily between E-P pairs, making up 58.8% of the total loops. This is consistent with the known associations of these histone marks to promoter and enhancer CREs^63–65^. To allow more quantitative analyses of these large-scale chromatin interactions we transformed the resulting HiChIP looping data into a graphical network with each node representing a CRE and the edges being the chromatin loops between these two elements (**Fig 1D**). With this regulatory network, we then overlaid the various multi-omics datasets to provide a framework that can interrogate the impact of AR binding. We investigated androgen-driven AR-mediated gene transcription based on the previously characterized hallmark androgen-responsive genes (**Supp Fig 1A, B**). As expected, these were induced compared to similarly expressed random AR-independent genes (**Fig 1E**). This comprehensive multi-omics dataset and graphical regulatory network provided a framework to quantitatively investigate the temporal impact of AR binding on epigenetic features, chromatin looping, and gene expression.

### Androgen stimulation leads to stepwise epigenetic changes

With this structured regulatory network, we then explored how AR binding affects each epigenetic feature and its relationship with gene transcription. The AR-regulated genes’ promoters (P+AR) and their looped AR-bound enhancers (E+AR) were compared to randomly matched from highly expressed AR-independent genes’ promoters (P-AR) and their looped AR-free enhancer (E-AR) (**Fig 2A**). We observed that E+AR displayed a strong AR and FOXA1 signal following androgen stimulation, that reached its peak at 4h and then subsequently decreased at 16h and 72h. Emphasizing FOXA1’s pioneering activity, E+AR displayed elevated FOXA1 signal at the initial time point (0m). As expected, the AR and FOXA1 signal did not significantly change at AR-independent promoters or AR-free enhancer. Interestingly, those FOXA1-bound CREs that were not co-occupied by AR had no change in the relative FOXA1 signal (**Supp Fig 1C, D**), suggesting that AR potentially influences FOXA1 occupancy^66^. As expected, we observed a higher H3K4me3 ChIP-seq signal at promoters compared to enhancers. This signal was largely unaffected by AR binding, but there were selective genes, including *KLK3*, which exhibited an increasing H3K4me3 mark at its promoter following androgen treatment (**Supp Fig 2**). We observed an elevated H3K27ac signal at promoters compared to enhancers (**Fig 2A**). Further, the H3K27ac signal increased specifically at E+AR, while it remained unchanged at E-AR. This change at E+AR was also observed for chromatin accessibility (ATAC-seq), though the maximum signal (16h) was found to occur after AR and FOXA1 peak occupancy (4h). We also investigated the epigenetic changes of AR down-regulated gene promoters and their AR-bound enhancers (**Supp Fig 3A, B)**. Interestingly, we observed that the AR-bound CREs (E+dAR), that were looped to the promoter of down-regulated genes (P+dAR) displayed a very similar increase in AR, FOXA1, H3K27ac, and chromatin accessibility (**Fig 2B**). There was no statistically significant change at any time point in the AR-bound enhancers of either up or downregulated genes (*p>0.1*). Overall, these kinetic datasets show that there is a sequential process that occurs following androgen stimulation where AR, FOXA1, and H3K27ac signals selectively increase at looped enhancers before inducing subsequent changes in chromatin accessibility.

### AR-bound enhancers increase contact frequency to AR-upregulated gene promoters

We first investigated how chromatin looping changed following AR activation. Initially, we compared the number of loops formed following androgen stimulation. We found that both promoters of AR-regulated genes (P+AR) and their looped AR-bound enhancers (E+AR) did not exhibit any significant changes (*p*>0.05) in the number of loops compared to background P-AR/E-AR (**Fig 3A**). Supporting this, there was no significant difference in the distribution of the number of loops on gene promoters during androgen treatment (**Supp Fig 4A**). These results demonstrate that AR binding does not cause a substantial rewiring of chromatin looping.

While androgen treatment did not significantly change the number of loops, we did observe an increase in contact frequency at AR-bound enhancers looped to AR-regulated genes (**Supp Fig 5**). To quantify these changes, we calculated the fold change in contact frequency compared to bootstrapped (*b*=1000) random AR-independent genes from both promoter and enhancer viewpoints (**Fig 3B**). We observed that AR activation increased the contact frequency of loops at AR-regulated promoters (P+AR) over time, with a peak at 16h (H3K27ac: *p*<0.001; H3K4me3: *p*<0.001) (**Fig 3C**). In contrast, the change in contact frequency of loops to promoters of AR independent genes (P-AR) remained stable. From an enhancer viewpoint (**Fig 3D**), AR-bound enhancer CREs (E+AR) showed a significant increase in both H3K27ac and H3K4me3 contact frequency (H3K27ac: *p*<0.001; H3K4me3: *p*<0.001). In contrast, AR-free CREs (E-AR; grey) that were connected to the same AR-regulated gene promoters did not exhibit any significant change. Further, no change in contact frequency was observed in the CREs (E-ARi; black) that interact with AR-independent gene promoters. While chromatin contact frequency increased between AR-bound enhancer CREs (E+AR) and upregulated hallmark gene promoters (P+AR), we did not observe any change in loops between AR-bound CREs (E+dAR) and downregulated gene promoters (**Fig 3E**). This is particularly striking as, AR-bound CREs (E+AR, E+dAR) exhibited a similar pattern in their epigenetic profiles, regardless of whether they looped to an upregulated or downregulated gene (**Fig 2A**, **Fig 2B**). Given that a similar trend is observed at all AR-bound CREs, this suggests that epigenetic modifications must be combined with chromatin looping to contribute to gene expression.

To interrogate the kinetic changes in epigenetics and contact frequency we focused on several up-regulated androgen-responsive genes (*KLK2*, *KLK3*, *NKX3-1*, *UAP1*, *ABCC4*, and *DHCR24*) (**Fig 3F**). As expected, AR-free enhancers (E-AR; grey) had minimal changes in epigenetics and contact frequency. Interestingly, we observed that AR-bound enhancers (E+AR; orange) had broad heterogeneity in response to treatment. The same enhancer typically had both the highest change in epigenetic features and contact frequency. Our findings demonstrate that AR-bound enhancers increase contact frequency with AR-regulated gene promoters in response to androgen treatment. However, this response is heterogeneous and there is significant variability among AR-bound enhancers.

### Association of nascent transcription to epigenetic changes and contact frequency

To determine if the change in contact frequency precedes or occurs simultaneously with AR-mediated gene transcription, we characterized the kinetic rate of androgen-induced gene expression. We identified up-regulated androgen-induced genes from nascent RNA (Start-seq) and categorized them based on the maximal expression (30m, 4h, 16h or 72h) (**Fig 4A**). We chose to group these genes based on nascent RNA, as RNA-seq provides only steady-state quantification of mRNA transcripts^67^. Between these groups, no significant difference was observed in the maximal signal at AR-bound enhancers for AR, FOXA1 and H3K27ac ChIP-seq or chromatin accessibility (**Fig 4B**), suggesting that the epigenetic features do not dictate the timing of maximal nascent RNA production. However, we observed a temporal relationship between nascent transcription and chromatin contact frequency. Specifically, the maximal nascent transcription (4h and 16h groups) occurred simultaneously with an increase in H3K27ac and came before the peak of H3K4me3 (16h) maximal contact frequency at AR-bound enhancers (**Fig 4C**). These observations indicate that the change in contact frequency does not precede RNA transcription. Overall, these findings suggest that although AR and FOXA1 rapidly bind at enhancers upon androgen treatment, the maximal transcription occurs either simultaneously with or before a maximal alteration in chromatin contact frequency.

### Multi-enhancers influence transcription proportional to contact frequency

Having observed marked heterogeneity in chromatin loop contact frequency at AR-bound enhancers, we began to explore the impact of multi-enhancer contacts on gene expression. We expand the scale of our analysis across all detectable genes (n=13,575), and characterized how loop frequency correlates to gene expression. Most genes had multiple CREs interacting with target promoters with a median frequency of ∼15 interactions per gene (**Supp.** Fig 6A). With this large dataset, we evaluated three models correlating H3K27ac and H3K4me3 contact frequency to gene expression where: all interactions contribute equally to gene expression (average), only a single strong interaction affects gene expression (maximum) and all interactions contribute to gene expression in an additive manner (sum) (**Fig 5A**). While all models strongly correlated H3K27ac and H3K4me3 contact frequency to gene expression (*p* < 0.0001; Spearman’s test), we found that the sum model (H3K27ac; r=0.896, H3K4me3; r=0.904) and maximum model (H3K27ac; r=0.839, H3K4me3; r=0.876) correlated significantly better than the average model (H3K27ac; r=0.643, H3K4me3; r=0.679) across all time points (**Fig 5B and Supp Fig 7**). To quantify the importance of each feature, we built a random forest regressor to predict the expression from these operations (Accuracy; r^2^=0.805) and found that the additive model (sum) best-predicted expression (**Supp Fig 6B**)^68^. However, as we observed similar correlation with the maximum model, which scored only a single chromatin loop, this suggested that the contact frequency of chromatin loops to a promoter were markedly unbalanced and that there was a “dominant” interaction. To characterize this, we quantified the inequality of chromatin loop contact frequency to a target promoter and found that both H3K27ac and H3K4me3 were strongly unbalanced (*Gini* >∼ 0.5; **Supp Fig 6C**). These findings suggest that enhancers interacting with gene promoters do not have a uniform distribution in contact frequency, and that there are “dominant” loops that strongly correlate with expression (**Fig 5D**).

To better understand how these potential dominant loops are dynamically affected by acute perturbation, we characterized the CRE-promoter interactions of androgen-regulated genes (n=88; **Supp Table 2**). Dominant loops were identified for every gene promoter by first scaling the contact frequency of interacting CREs in the range (0, 1), and selecting the highest ones with a threshold (0.8). The dominant loops were not solely based on proximity to the gene promoter as they were the closest CRE in only 27% of AR-regulated genes (n=24) at any time point. Suggesting that the dominant loops are “primed” before AR binds, the same dominant loops were commonly found at every time point (**Supp Fig 8**). Furthermore, we found that the dominant AR-bound CREs (D+) had significantly (p<0.0001) higher dynamic change in contact frequency than non-dominant AR-bound CREs (D-) that interacts with AR-regulated gene promoters (**Fig 5C**). Interestingly, the dominant loops were highly gene specific. When we characterized two AR-regulated genes, *KLK3* and *KLK2*, that share many looped CREs (**Fig 5E**), we observed gene-specific disparities in contact frequency changes from an AR enhancer (ARBS3) to either the *KLK2* or *KLK3* promoters. While ARBS3 displayed the strongest and most dynamic contact frequency with the *KLK2* promoter, it did not exhibit the strongest change for the *KLK3* promoter. Instead, the loop with well-known upstream AR-enhancer^69^ (ARBS2) dominates others. Interestingly, both dominant loop enhancers were not the CRE with the highest change in AR peak height, suggesting that there are additional factors that contribute to changes in contact frequency. Overall, these results demonstrate that AR binding does not affect chromatin looping equally and that those CREs with the most dynamic contact frequency potentially have a greater impact on gene transcription.

### CRISPR-based perturbations confirm the existence of “dominant” chromatin loops

To experimentally validate these correlative models, we tested the effects of dominant chromatin loops on androgen-induced gene expression. Utilizing CRISPRi, a derivative of the CRISPR/Cas9 system that suppresses enhancer activity without altering DNA sequences, we inhibited all AR-bound enhancers (n=20) that interact with (**Supp Table 1, Supp Fig 9**) the previously described AR-regulated genes (*KLK2*, *KLK3*, *NKX3-1*, *UAP1*, *ABCC4*, and *DHCR24*). Across all tested genes we found that inhibiting those CREs, that had a dominant chromatin loop, significantly impacted androgen-induced transcription (**Fig 6A**). Importantly, as we observed highest dynamic change in contact frequency at ARBS3 and ARBS2 for KLK2 and KLK3, respectively (**Fig 5E**), the highest gene expression inhibition is observed at these ARBSs. Supporting the kinetics of the dominance model, we observed an inverse correlation between the gene expression inhibition and the dynamic change in contact frequency of loops during androgen stimulation (**Fig 6B**). This correlation was greater than other genomic features including AR, FOXA1, H3K4me3, H3K27ac and chromatin accessibility (**Supp Fig 10**). These results demonstrate that not all CREs contribute equally to androgen stimulation and that their contact frequency to promoters correlates with their impact on gene expression. Overall, these results underscore the importance of spatial genome organization in transcriptional regulation and validate our proposed multi-enhancer dominance model.

**Figure 6:**
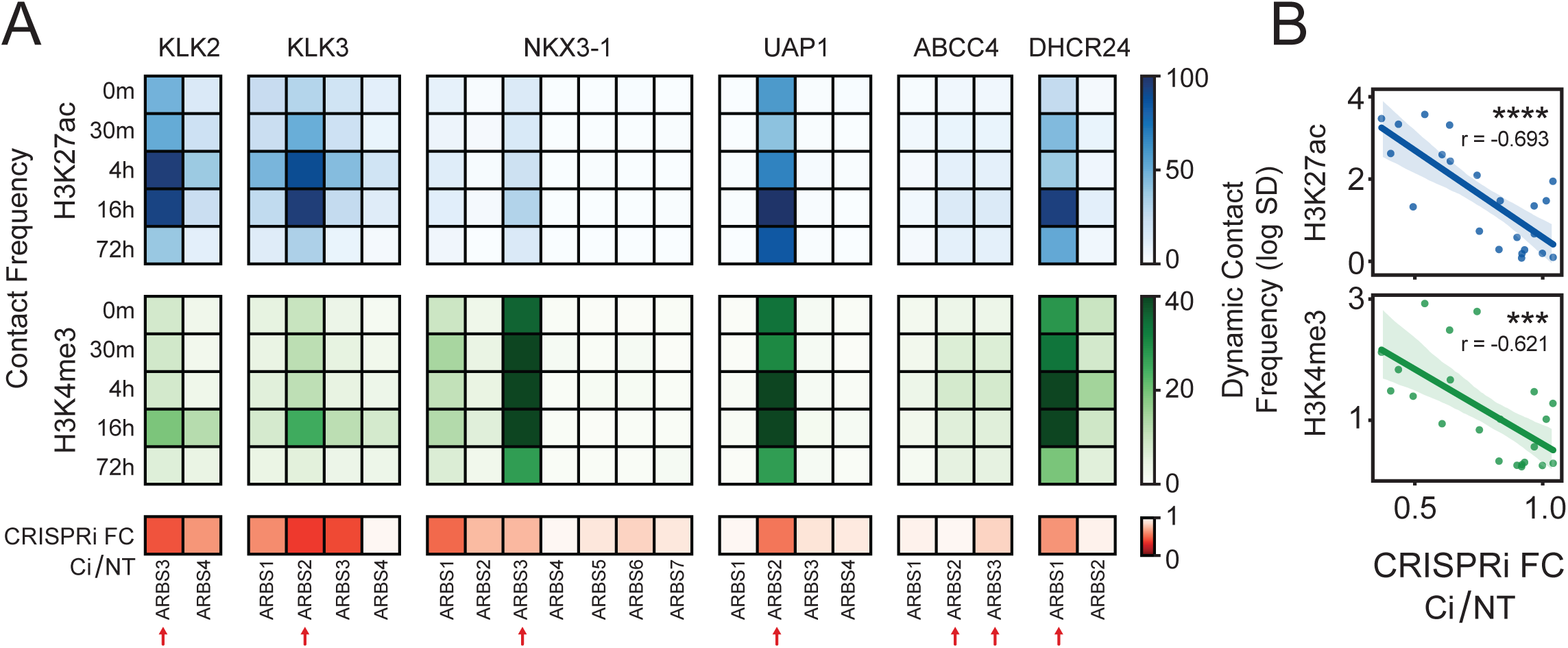
Androgen induced gene expression is significantly affected by perturbing the most dominant AR-bound enhancers. **(A)** Functional characterization of individual AR-bound CRE on androgen-induced expression. Heatmap of contact frequency strength at each time point (H3K27ac: blue, H3K4me3: green) for individual AR-bound CREs to a target gene promoter (*KLK2, KLK3, NKX3-1, UAP1, ABCC4* and *DHCR24*). Each AR-binding site (ARBS) was individually inhibited with CRISPRi and treated with androgen (10nM DHT). The inhibition of the target gene induction was calculated compared to the non-targeting control (CRISPRi FC Ci/NT; red). Identified dominant enhancers according to contact frequency are represented with red arrow. **(B)** Correlation of CRISPRi induced gene-knockdown and chromatin loop dynamic (Standard Deviation in time) contact frequency for each ARBS. For all data ns p>0.05, * p<0.05, ** p<0.01, *** p<0.001, **** p<0.0001.

## DISCUSSION

Transcription factors bind to specific DNA sites and regulate gene expression through the recruitment of co-regulatory proteins that activate transcriptional machinery^70,71^. Yet despite extensive research, many questions remain about how this complex process occurs, particularly as there are multiple CREs that interact with each promoter. Using the ligand-activated AR as a model system, we characterized how transcription factor activation changes the epigenetics and chromatin looping. Similar to published studies, our work showed that the AR stabilizes/recruits FOXA1 and increases both H3K27ac and chromatin accessibility (**Fig 2A**)^2,5,7,72,73^. However, by characterizing a kinetic dataset, we showed that the change in chromatin accessibility occurs after both FOXA1/AR binding and H3K27ac post-translational modifications, suggesting that this is mediated by the recruitment of additional chromatin-remodeling proteins. Those enhancers that are not bound by AR do not exhibit these changes. However, these epigenetic changes do not solely drive gene expression, as we observed a consistent pattern in all AR-bound CREs, regardless of either the direction of expression (upregulated and downregulated), or timing of maximal nascent transcription (**Fig 2**; **Fig 4B**). These observations suggest a sequential mechanism, independent of gene expression, in which AR activation recruits specific co-regulatory proteins that alter histone modification and chromatin accessibility.

This work also characterized how AR binding affects genome organization. This is a controversial field, with earlier studies proposing that steroid hormone receptors significantly reorganize chromatin looping when activated^74^. However, recent work has suggested that gene expression may occur through already-existing interactions^23^. Our research found that androgen treatment does not cause rewiring of chromatin contact loops but instead increases the contact frequencies of previously established loops (**Fig 3C, D; Supp Fig 5**). Interestingly, we observed that maximal chromatin looping happens either during or after the maximal nascent gene expression (**Fig 4C**). This suggests that increasing chromatin looping does not precede, but instead likely occurs simultaneously with gene transcription. Supporting this result, recent work observed that higher nascent RNA production is associated with a higher frequency of chromatin contacts^22^. Changes in the chromatin contact frequency have also been shown to be associated with differential gene expression^20,23^. Although several studies observed temporal changes in chromatin looping that occur before maximal RNA expression^21,23^, this distinction is likely due to the experimental methodology used, as RNA-seq primarily quantifies mature mRNA, whereas Start-seq captures only nascent mRNA. Highlighting the consistent pattern of epigenetic features of AR-bound enhancers (**Fig 2**; **Fig 4B**), we can infer chromatin looping is an additional mechanism which regulates gene expression independently of AR binding. The importance of chromatin loop contact frequency is highlighted when we look at AR-downregulated genes, which show similar changes in epigenetic alterations and chromatin accessibility but not contact frequency (**Fig 2B**; **Fig 3E**). This suggests that AR binding recruits additional factors that potentially increase the contact frequencies of pre-established loops between enhancers and their target gene promoters to regulate the gene expression.

Numerous studies demonstrate that multiple enhancers contribute to the expression of a single gene^7,75^. However, there is no consensus about how each CRE contributes to gene expression. Our work suggests that connected enhancers contribute unevenly, and there exists dominant loops that have the largest impact on gene expression. These findings align with the assumptions made in the activity-by-contact (ABC) model^76^, which presumes that the enhancer’s impact on gene expression is correlated with the strength of the contact between them. Both our validation (**Fig 6**), and a recent large-scale CRISPRi study from Barshard *et al.*^22^, demonstrate that those enhancers with higher contact frequency to target promoters are more likely to be functionally important which suggest a “dominance” model (**Fig 5D**). However, further improvements of this model can help us to understand the effect of individual CREs, thus allowing us to evaluate how multiple transcription factors impact gene expression.

Given the complex nature of this system, we had to make several assumptions. First, we chose to define individual regulatory units based on chromatin accessibility, since defining enhancers and promoters by histone modifications is prone to false positives^77,78^. Within these CREs, we focused on hallmark androgen response genes (**Fig 1E**), as these have been extensively demonstrated to be regulated by AR. Next, we used HiChIP, a protein-centric HiC method, rather than conventional HiC to provide enhanced resolution and specificity in capturing protein-bound chromatin^79^. By characterizing both promoter-centric (H3K4me3) and enhancer-centric (H3K27ac) chromatin loops, we believe this provided us with different genomic perspectives of chromatin loops and reduced potential biases (**Fig 1B, C**)^21,80,81^. It is important to note that while H3K27ac is strongly impacted by AR binding we observed that H3K4me3 was relatively static (**Fig 2**; **Fig 4B**), suggesting that the change in contact frequency (**Fig 3C-E**; **Fig 4C**) following androgen treatment was not due to capture efficiency. Finally, as gene expression regulation circuits are challenging to integrate due to the intricate network structures formed by diverse CREs (**Fig 1D**), we utilized regulatory networks wherein the nodes and edges represent the CREs and chromatin loops, respectively. Several studies have employed graph-based approaches to connect genomic regions and address various high-dimensional biological inquiries^82–84^. The versatility of this approach underscores the quantitative advantages of utilizing graphs to characterize chromatin loops. Despite these limitations, this study represents one of the largest experimental datasets (**Fig 1A**) to characterize AR-mediated transcription.

Overall, this paper provides new insight into several important aspects of AR-mediated gene expression. We show that AR binding triggers a temporal cascade which increases FOXA1 and H3K27ac that affects chromatin accessibility. Further, we demonstrate that AR does not introduce novel chromatin loops, but instead increases the contact frequency between AR-bound enhancers and their target promoters. However, the effect of each enhancer on gene expression is markedly heterogeneous and proportional to promoter contact frequency. These findings suggest that while AR binding to DNA induces a step-wise epigenetic alteration, the impact of bound enhancers is strongly dependent on the contact frequency of the established chromatin loops with the target promoter.

## ACKNOWLEDGMENTS

This work was supported by funding from TUBITAK (221Z116).

---

**Supplementary Figure 1:**
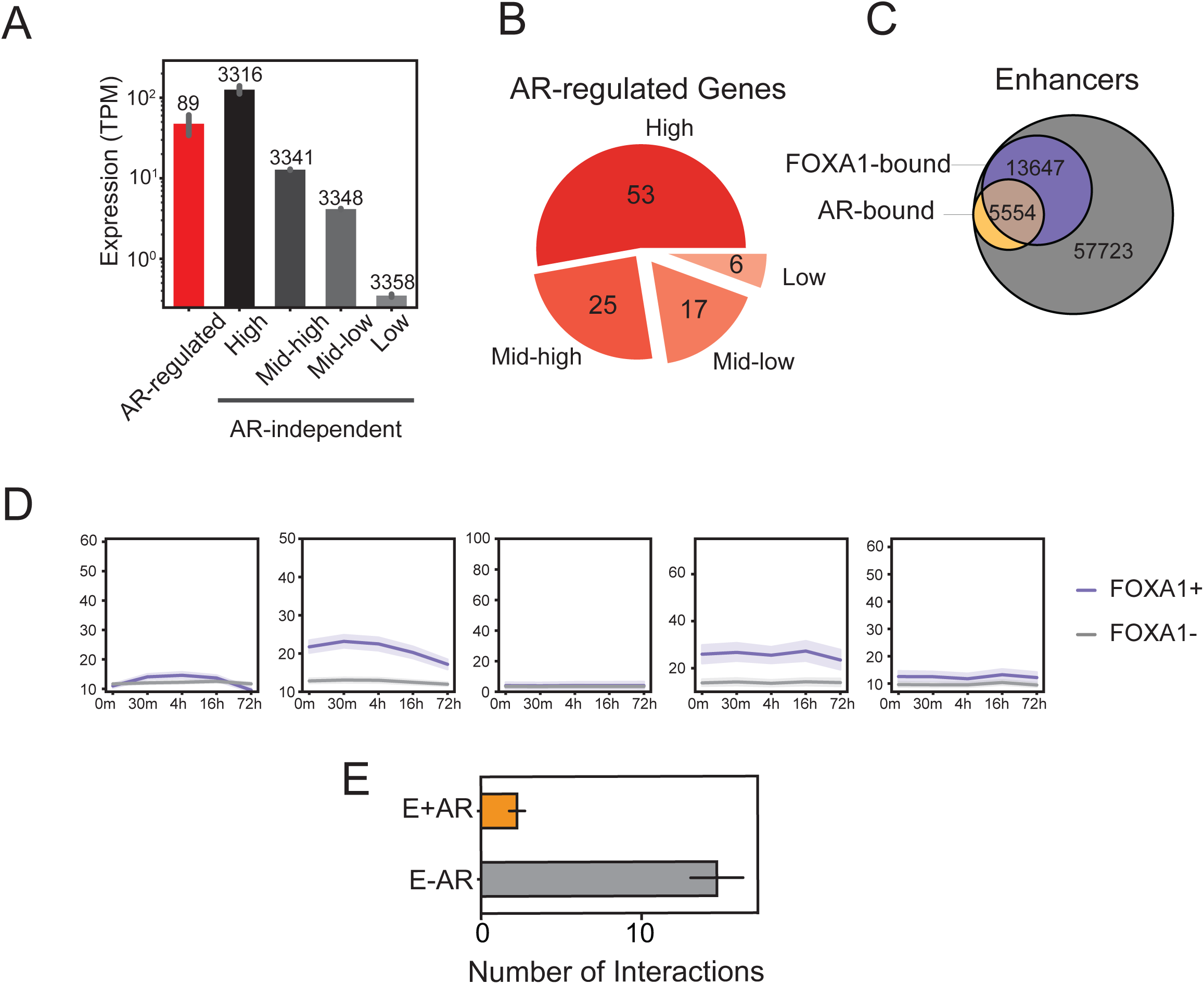
We split the transcriptome into four equal-size groups according to their average expresion. **(A)** The average expression of AR-regulated genes (red) and AR-independent gene quartiles (greys). **(B)** Pie chart illustrating the distribution of AR-regulated genes among assigned quartiles. **(C)** Venn diagram representing AR-bound, FOXA1-bound enhancers. **(D)** Epig-enome changes (see Fig 2) of AR-free enhancers that are either FOXA1-bound (purple) or FOXA1-free (grey). **(E)** Number of E+AR and E-AR loops of AR-regulated genes.

**Supplementary Figure 2:**
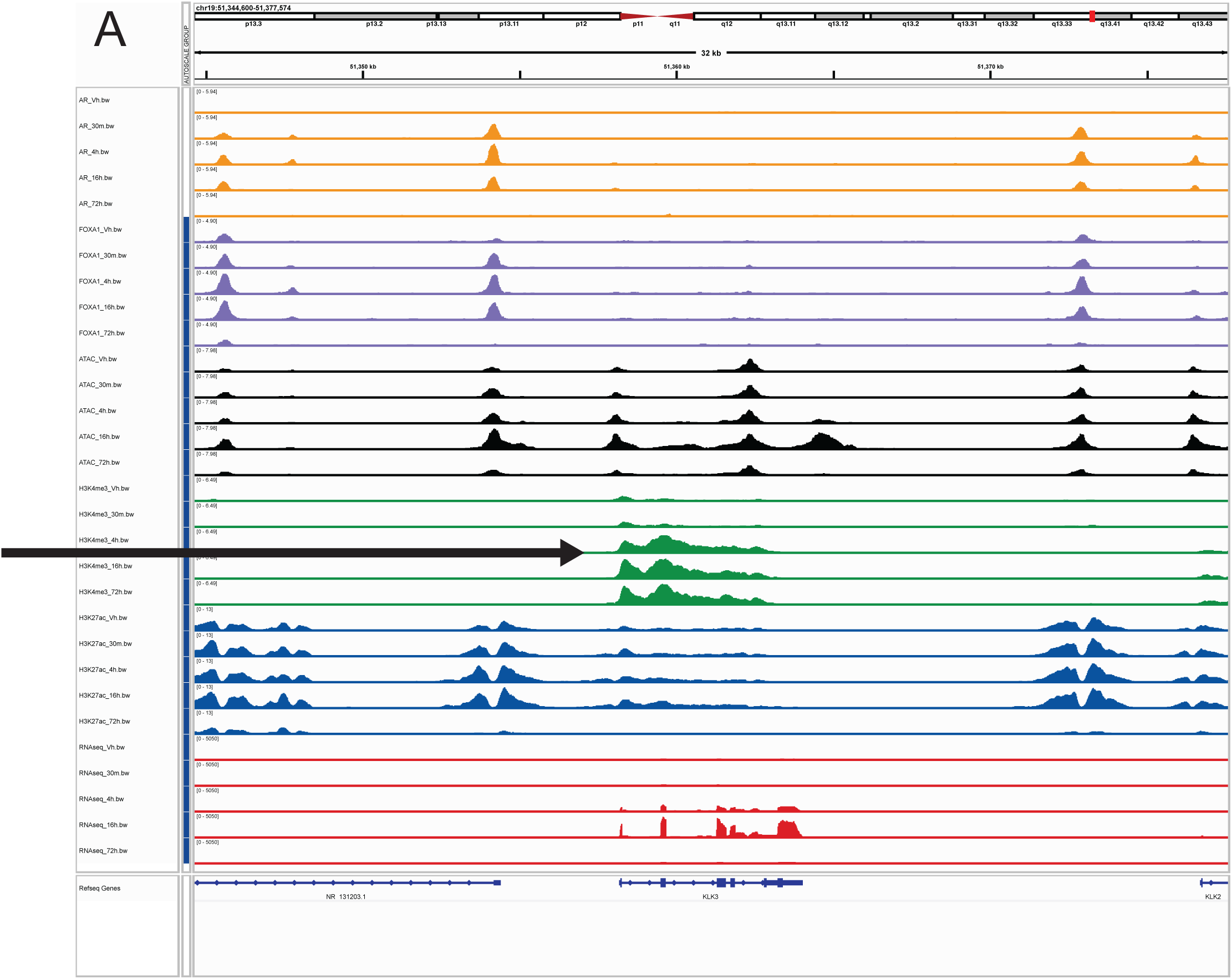
IGV track for KLK3 (∼15kb) gene locus representing AR (yellow), H3K4me3 (green), H3K27ac (blue), RNA-seq (red) tracks. The arrow represents the H3K4me3 signal at 4h on the KLK3 promoter.

**Supplementary Figure 3:**
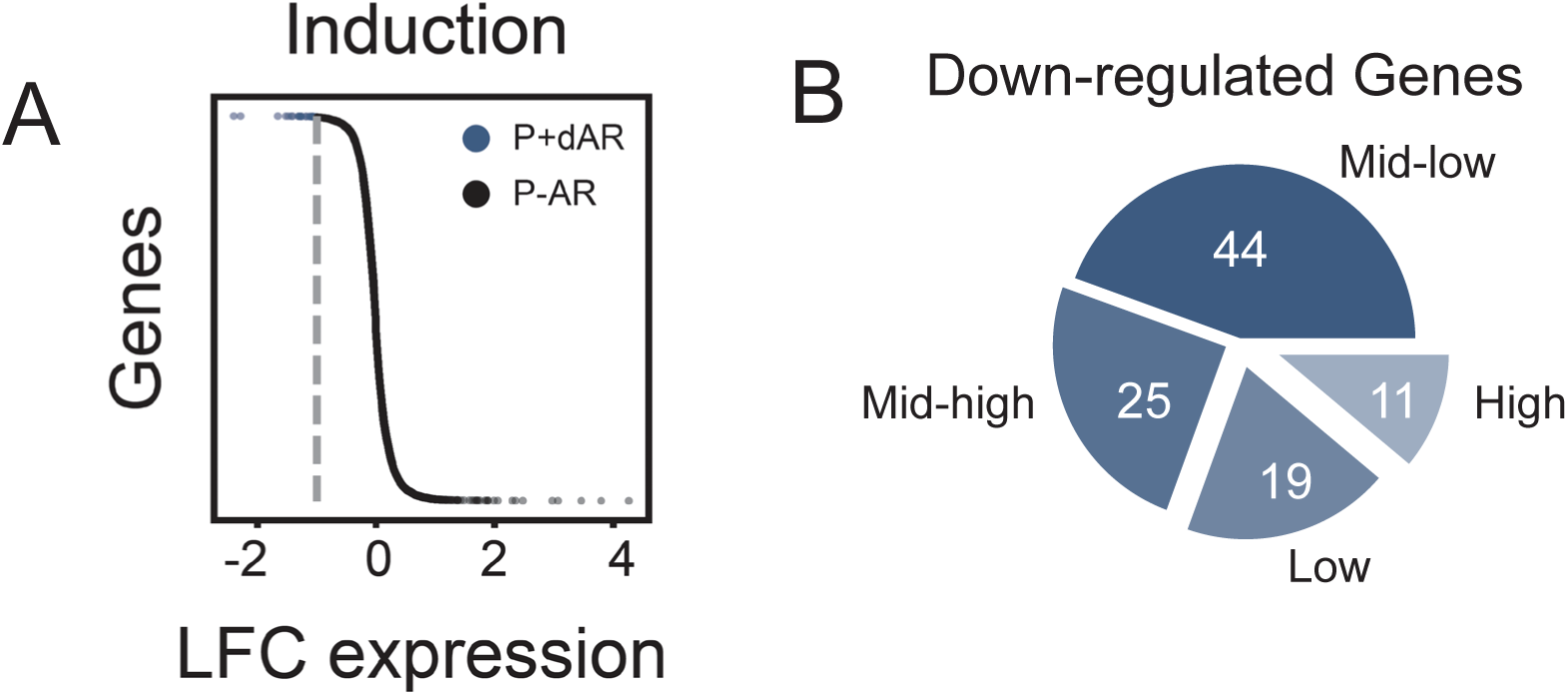
Identification of down regulated genes. **(A)** The downregulated genes are selected according to 16h vs 0m Log2FoldChange (-1 >). **(B)** The number of genes in each quartile.

**Supplementary Figure 4:**
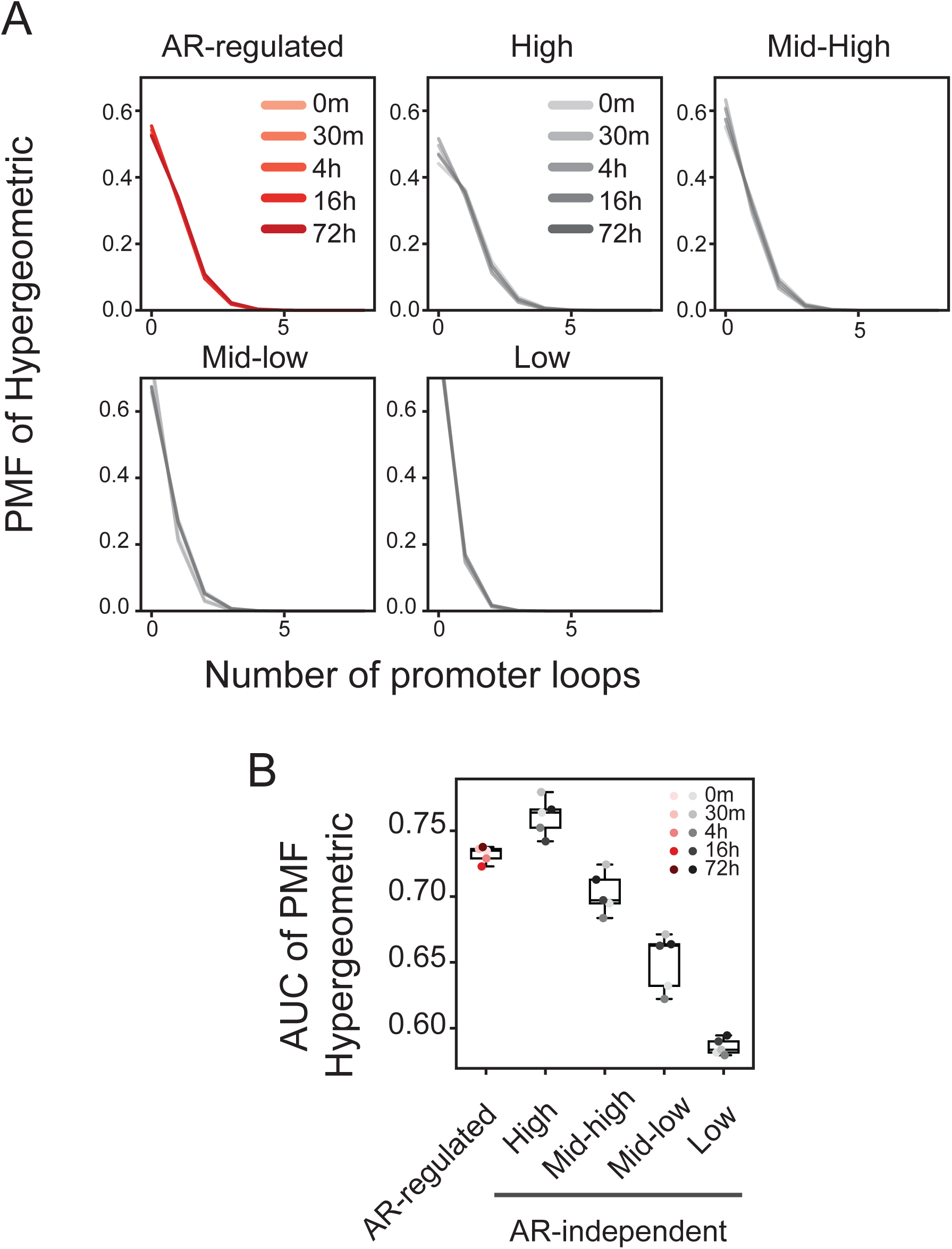
**(A)** Probability mass function of hypergeometric distribution of the number of loops on gene promoters for each time point. Each pane represents one gene set (AR-regulated or expression quartiles). **(B)** The area-under-curve (AUC) of the PMF of hypergeometric (see A).

**Supplementary Figure 5:**
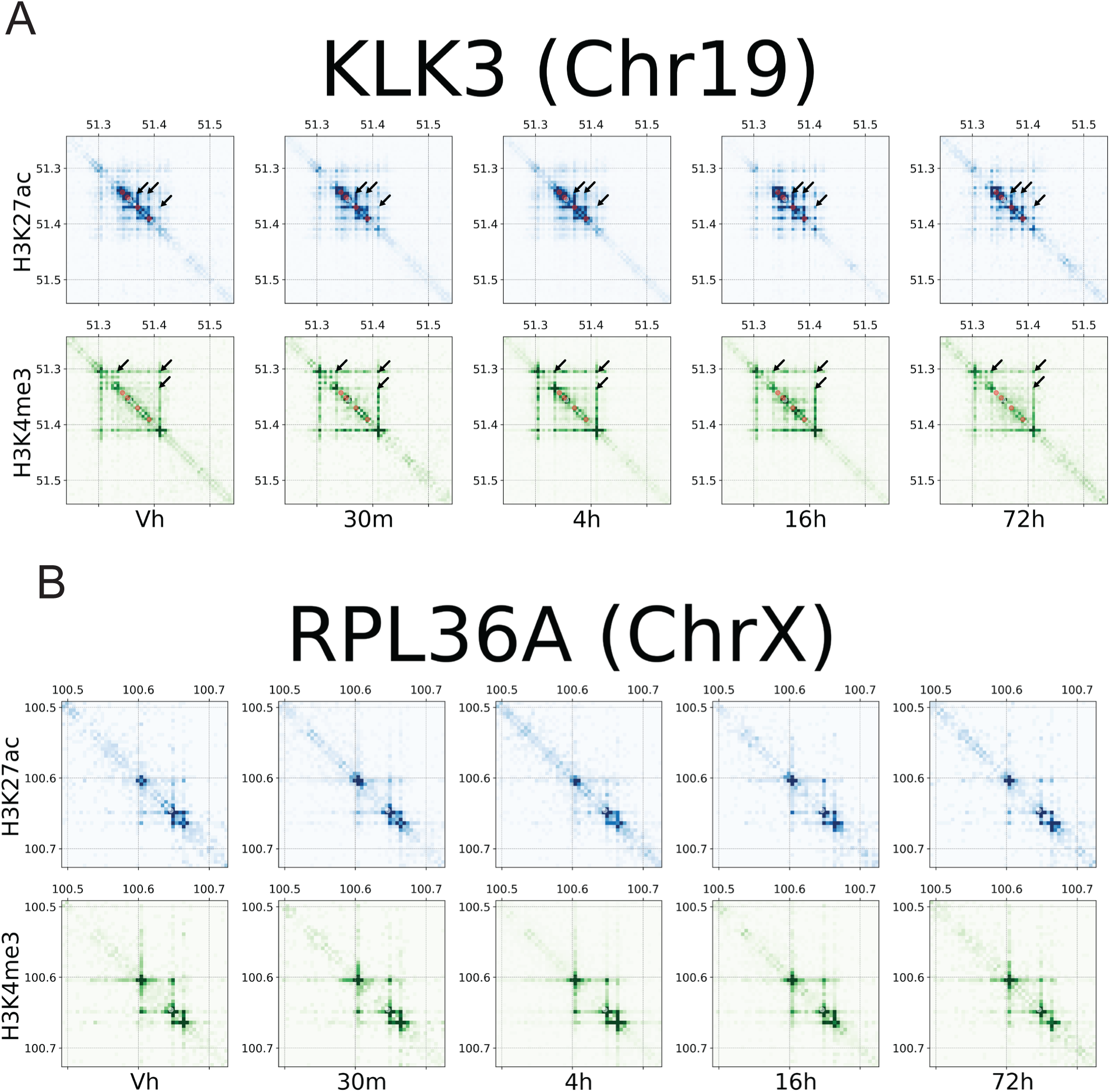
Contact maps of H3K27ac (top) and H3K4me3 (bottom) centering the gene promoters (white) and their putative AR-bound enhancers. +/-150kb with 5kb resolution **(A)** KLK3 locus, an androgen responsive gene. The arrows represent increasing contact frequency of AR-bound enhancer loops. **(B)** RPL36A locus, a highly expressed gene that has no AR-bound enhancers.

**Supplementary Figure 6:**
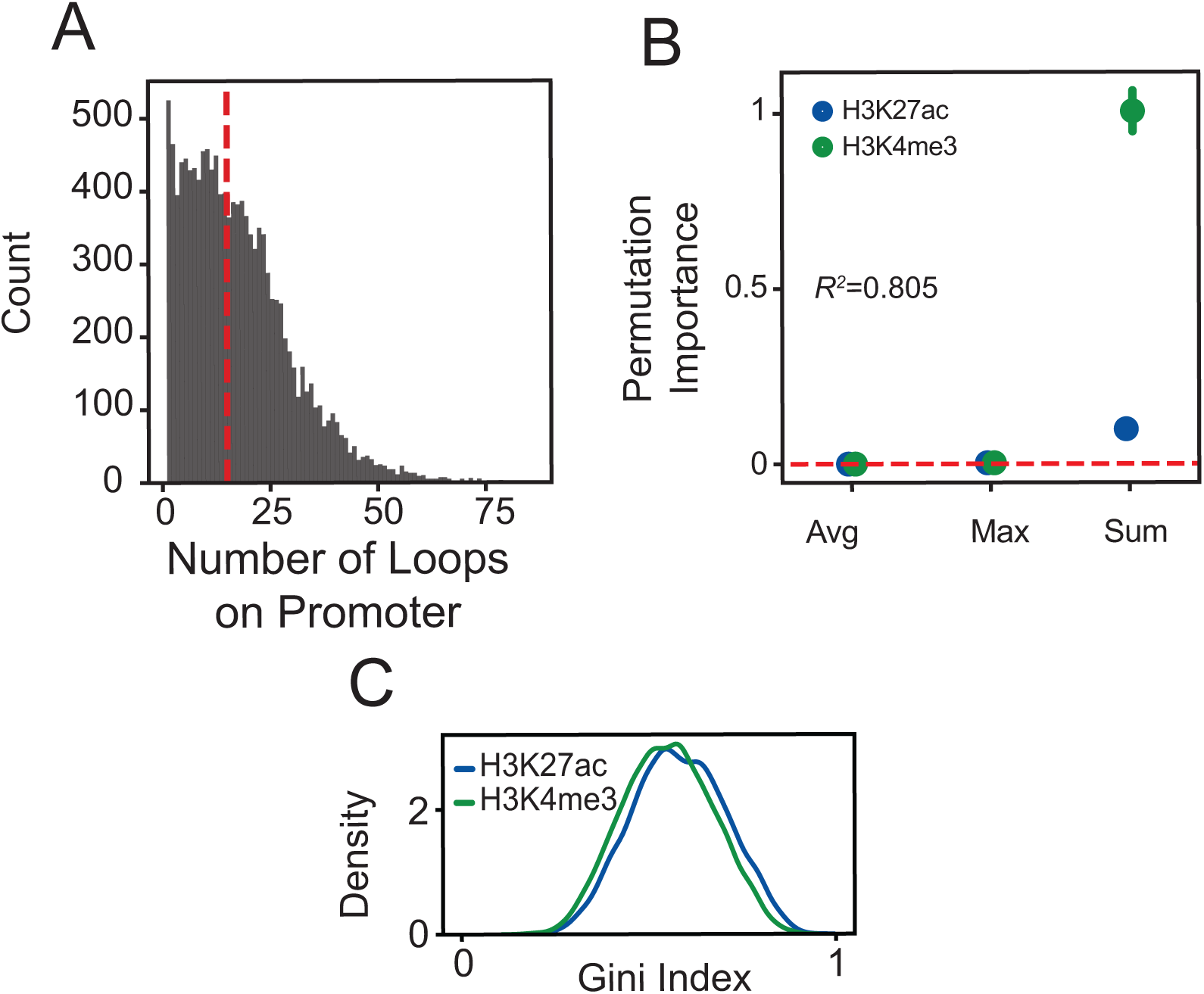
**(A)** Number of loops on gene promoters. **(B)** Random forest regressor feature permutation importance. **(C)** Density plot representing Gini-index for AR-regulated (P+AR) and AR-independent (P-AR) gene promoters.

**Supplementary Figure 7:**
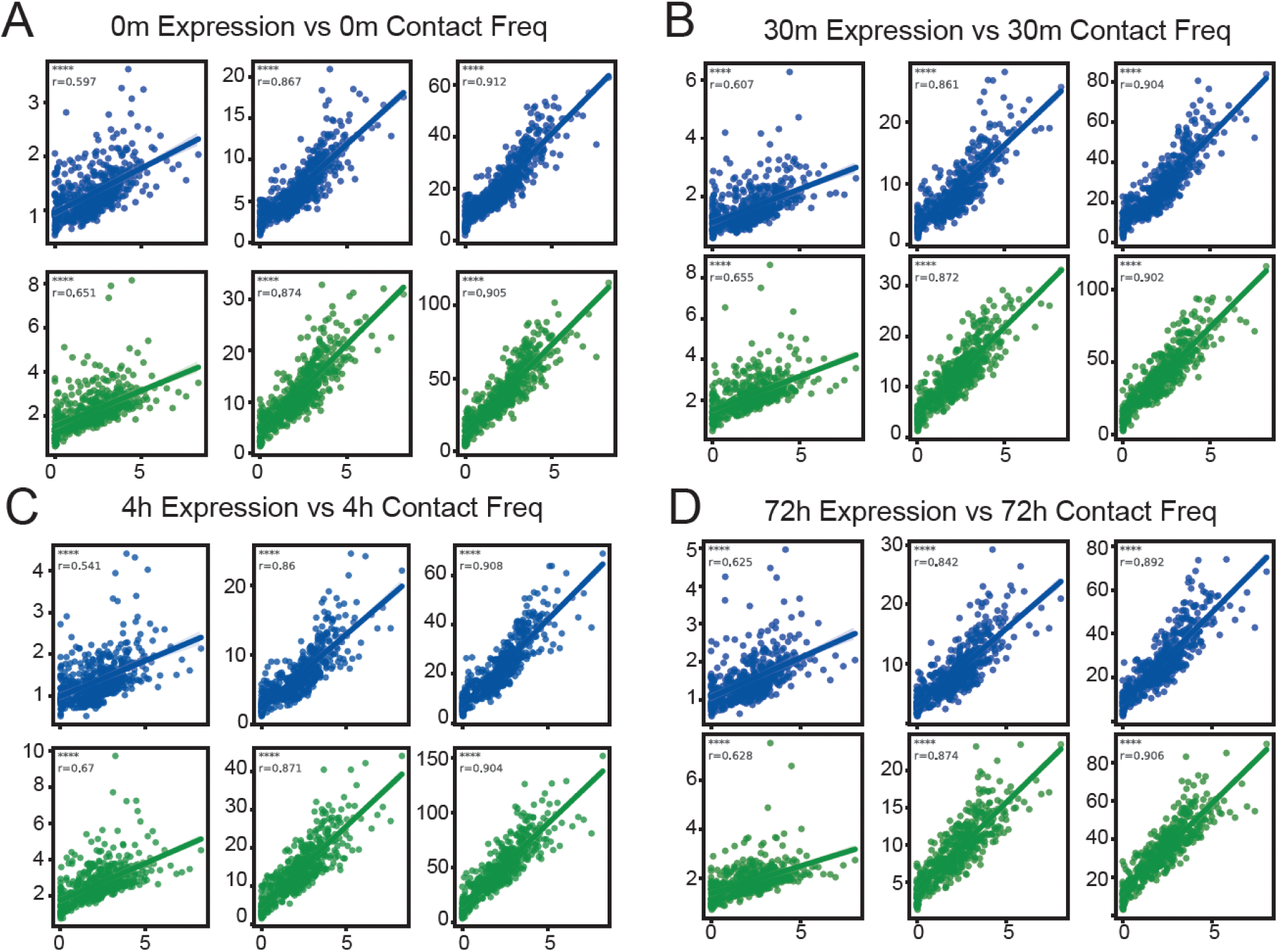
Binned (k=25) scatter plot of log expression (x-axis) vs. contact frequency (y-axis) according to the summarization function. Columns are avg, max, sum respectively. The blue and green scatterplots represent H3K27ac and H3K4me3 **(A)** 0m **(B)** 30m **(C)** 4h **(D)** 72h data respectively. For all data ns p>0.05, * p<0.05, ** p<0.01, *** p<0.001, **** p<0.0001.

**Supplementary Figure 8:**
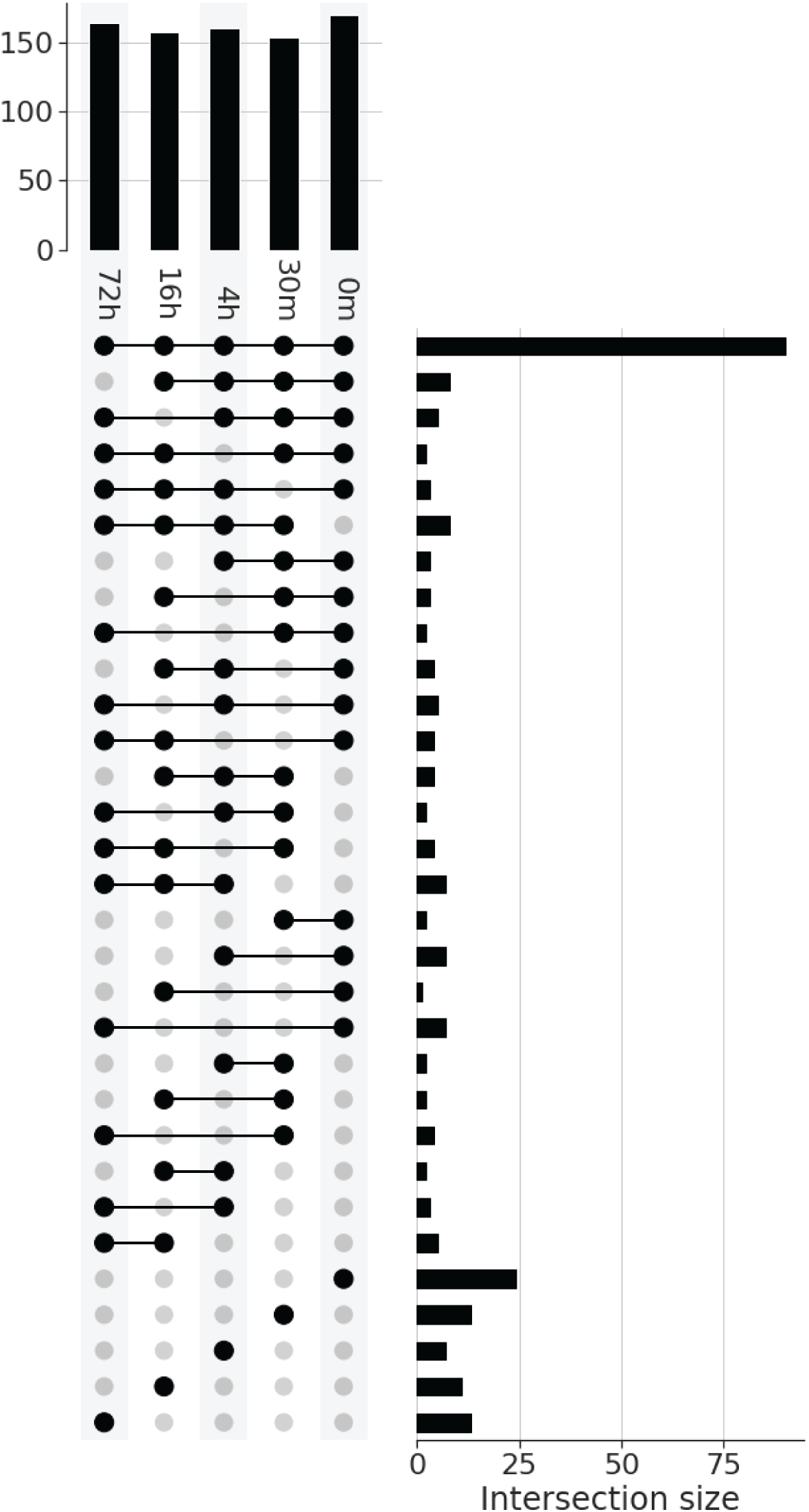
Dominant loops of AR-regulated genes are identified for every gene promoter by scaling contact frequency (H3K27ac or H3K4me3 separately) in range (0, 1), and selecting the highest ones with a threshold (0.8) at every time point (bar plot top). Most of the domintant loops are commonly found in every time point (upset plot).

**Supplementary Figure 9:**
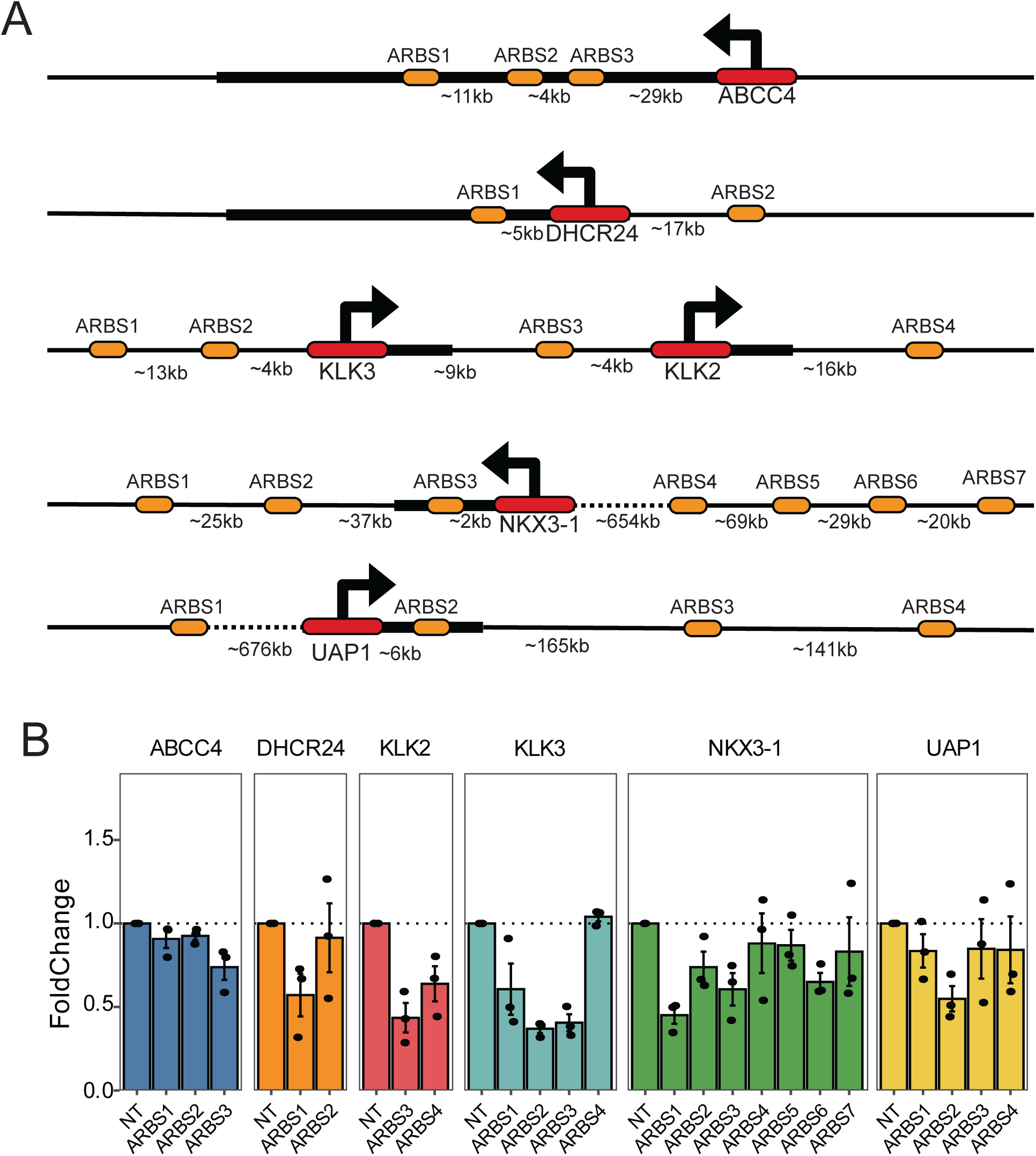
In total, 65 gRNA are desined for 20 ARBS (**Supp Table 1**). The experiment contains three biological replicates for each ARBS. **(A)** Genomic locations of each enhancer tested with respect to gene promoters’ location. **(B)** Gene expression changes in response to CRISPRi suppression of candidate enhancers.

**Supplementary Figure 10:**
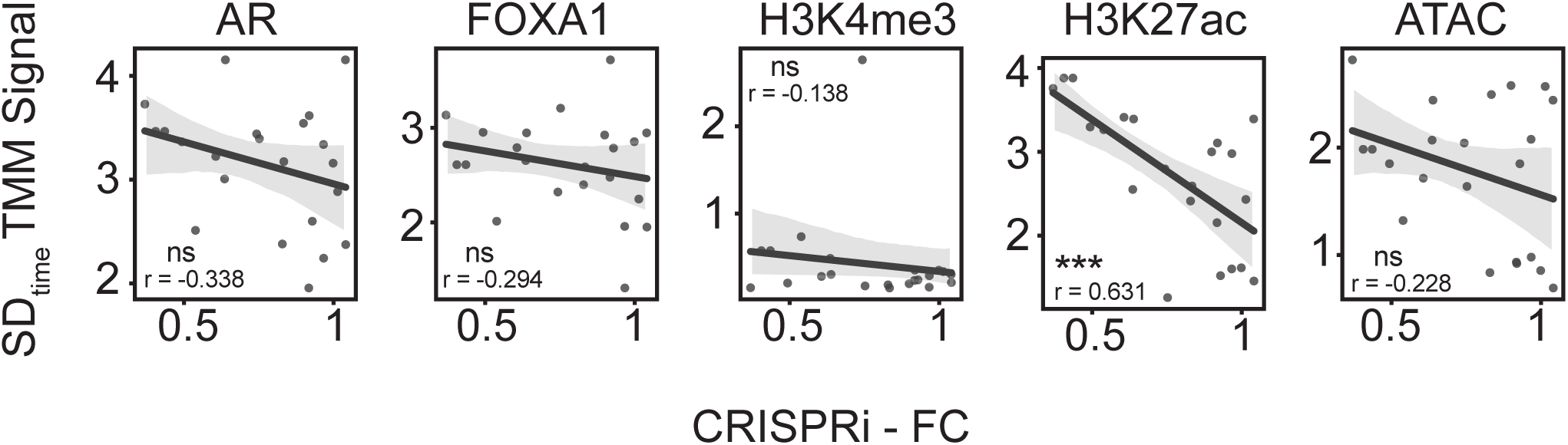
Changes in epigenetic features upon androgen treatment (SD_time_) vs. Gene expression changes in response to CRISPRi suppression of candidate enhancers.

## References

1. Nelson, P. S. et al. The program of androgen-responsive genes in neoplastic prostate epithelium. Proc. Natl. Acad. Sci. U. S. A. 99, 11890–11895 (2002).

2. Davey, R. A. & Grossmann, M. Androgen Receptor Structure, Function and Biology: From Bench to Bedside. Clin. Biochem. Rev. 37, 3–15 (2016).

3. Georget, V., Térouanne, B., Nicolas, J.-C. & Sultan, C. Mechanism of antiandrogen action: key role of hsp90 in conformational change and transcriptional activity of the androgen receptor. Biochemistry 41, 11824–11831 (2002).

4. Ozanne, D. M. et al. Androgen receptor nuclear translocation is facilitated by the f-actin cross-linking protein filamin. Mol. Endocrinol. 14, 1618–1626 (2000).

5. Teng, M., Zhou, S., Cai, C., Lupien, M. & He, H. H. Pioneer of prostate cancer: past, present and the future of FOXA1. Protein Cell 12, 29–38 (2021).

6. Sahu, B. et al. Dual role of FoxA1 in androgen receptor binding to chromatin, androgen signalling and prostate cancer. EMBO J. 30, 3962–3976 (2011).

7. Stelloo, S., Bergman, A. M. & Zwart, W. Androgen receptor enhancer usage and the chromatin regulatory landscape in human prostate cancers. Endocr. Relat. Cancer 26, R267–R285 (2019).

8. Wilson, S., Qi, J. & Filipp, F. V. Refinement of the androgen response element based on ChIP-Seq in androgen-insensitive and androgen-responsive prostate cancer cell lines. Sci. Rep. 6, 32611 (2016).

9. Heinlein, C. A. & Chang, C. Androgen receptor (AR) coregulators: an overview. Endocr. Rev. 23, 175–200 (2002).

10. Jin, H.-J., Kim, J. & Yu, J. Androgen receptor genomic regulation. Transl. Androl. Urol. 2, 157–177 (2013).

11. Kadauke, S. & Blobel, G. A. Chromatin loops in gene regulation. Biochim. Biophys. Acta 1789, 17–25 (2009).

12. Zhu, I., Song, W., Ovcharenko, I. & Landsman, D. A model of active transcription hubs that unifies the roles of active promoters and enhancers. Nucleic Acids Res. 49, 4493–4505 (2021).

13. Sahu, B. et al. Sequence determinants of human gene regulatory elements. Nat. Genet. 54, 283–294 (2022).

14. Morgan, M. A. J. & Shilatifard, A. Reevaluating the roles of histone-modifying enzymes and their associated chromatin modifications in transcriptional regulation. Nat. Genet. 52, 1271–1281 (2020).

15. Tsompana, M. & Buck, M. J. Chromatin accessibility: a window into the genome. Epigenetics Chromatin 7, 33 (2014).

16. Wang, H. et al. H3K4me3 regulates RNA polymerase II promoter-proximal pause-release. Nature 615, 339–348 (2023).

17. Tewari, A. K. et al. Chromatin accessibility reveals insights into androgen receptor activation and transcriptional specificity. Genome Biol. 13, R88 (2012).

18. Guo, H. et al. Androgen receptor and MYC equilibration centralizes on developmental super-enhancer. Nat. Commun. 12, 7308 (2021).

19. Sawada, T., Kanemoto, Y., Kurokawa, T. & Kato, S. The epigenetic function of androgen receptor in prostate cancer progression. Front Cell Dev Biol 11, 1083486 (2023).

20. Greenwald, W. W. et al. Subtle changes in chromatin loop contact propensity are associated with differential gene regulation and expression. Nat. Commun. 10, 1054 (2019).

21. Reed, K. S. M. et al. Temporal analysis suggests a reciprocal relationship between 3D chromatin structure and transcription. Cell Rep. 41, 111567 (2022).

22. Barshad, G. et al. RNA polymerase II dynamics shape enhancer–promoter interactions. Nat. Genet. 55, 1370–1380 (2023).

23. D’Ippolito, A. M. et al. Pre-established Chromatin Interactions Mediate the Genomic Response to Glucocorticoids. Cell Syst 7, 146–160.e7 (2018).

24. Stelloo, S. et al. Androgen receptor profiling predicts prostate cancer outcome. EMBO Mol. Med. 7, 1450–1464 (2015).

25. Pomerantz, M. M. et al. The androgen receptor cistrome is extensively reprogrammed in human prostate tumorigenesis. Nat. Genet. 47, 1346–1351 (2015).

26. Pomerantz, M. M. et al. Prostate cancer reactivates developmental epigenomic programs during metastatic progression. Nat. Genet. 52, 790–799 (2020).

27. Grbesa, I. et al. Reshaping of the androgen-driven chromatin landscape in normal prostate cells by early cancer drivers and effect on therapeutic sensitivity. Cell Rep. 36, 109625 (2021).

28. Huang, C.-C. F. et al. Functional mapping of androgen receptor enhancer activity. Genome Biol. 22, 149 (2021).

29. Staller, M. V. Transcription factors perform a 2-step search of the nucleus. Genetics 222, (2022).

30. Uyehara, C. M. & Apostolou, E. 3D enhancer-promoter interactions and multi-connected hubs: Organizational principles and functional roles. Cell Rep. 112068 (2023).

31. Carey, M. The enhanceosome and transcriptional synergy. Cell 92, 5–8 (1998).

32. Merika, M. & Thanos, D. Enhanceosomes. Curr. Opin. Genet. Dev. 11, 205–208 (2001).

33. Choi, J. et al. Evidence for additive and synergistic action of mammalian enhancers during cell fate determination. Elife 10, (2021).

34. Antosova, B. et al. The Gene Regulatory Network of Lens Induction Is Wired through Meis-Dependent Shadow Enhancers of Pax6. PLoS Genet. 12, e1006441 (2016).

35. Cannavò, E. et al. Shadow Enhancers Are Pervasive Features of Developmental Regulatory Networks. Curr. Biol. 26, 38–51 (2016).

36. Hong, J.-W., Hendrix, D. A. & Levine, M. S. Shadow enhancers as a source of evolutionary novelty. Science 321, 1314 (2008).

37. Kvon, E. Z., Waymack, R., Gad, M. & Wunderlich, Z. Enhancer redundancy in development and disease. Nat. Rev. Genet. 22, 324–336 (2021).

38. Baca, S. C. et al. Reprogramming of the FOXA1 cistrome in treatment-emergent neuroendocrine prostate cancer. Nat. Commun. 12, 1979 (2021).

39. Giambartolomei, C. et al. H3K27ac HiChIP in prostate cell lines identifies risk genes for prostate cancer susceptibility. Am. J. Hum. Genet. 108, 2284–2300 (2021).

40. Dorighi, K. M. et al. Mll3 and Mll4 facilitate enhancer RNA synthesis and transcription from promoters independently of H3K4 monomethylation. Mol. Cell 66, 568–576.e4 (2017).

41. Hsu, J. Y. et al. CRISPR-SURF: discovering regulatory elements by deconvolution of CRISPR tiling screen data. Nat. Methods 15, 992–993 (2018).

42. Bae, S., Park, J. & Kim, J.-S. Cas-OFFinder: a fast and versatile algorithm that searches for potential off-target sites of Cas9 RNA-guided endonucleases. Bioinformatics 30, 1473– 1475 (2014).

43. Dobin, A. et al. STAR: ultrafast universal RNA-seq aligner. Bioinformatics 29, 15–21 (2013).

44. Patro, R., Duggal, G., Love, M. I., Irizarry, R. A. & Kingsford, C. Salmon provides fast and bias-aware quantification of transcript expression. Nat. Methods 14, 417–419 (2017).

45. Frankish, A. et al. GENCODE 2021. Nucleic Acids Res. 49, D916–D923 (2021).

46. Cornwell, M. et al. VIPER: Visualization Pipeline for RNA-seq, a Snakemake workflow for efficient and complete RNA-seq analysis. BMC Bioinformatics 19, 135 (2018).

47. Quinlan, A. R. & Hall, I. M. BEDTools: a flexible suite of utilities for comparing genomic features. Bioinformatics 26, 841–842 (2010).

48. Qin, Q. et al. ChiLin: a comprehensive ChIP-seq and DNase-seq quality control and analysis pipeline. BMC Bioinformatics 17, 404 (2016).

49. Li, H. & Durbin, R. Fast and accurate short read alignment with Burrows–Wheeler transform. Bioinformatics 25, 1754–1760 (2009).

50. Zhang, Y. et al. Model-based analysis of ChIP-Seq (MACS). Genome Biol. 9, R137 (2008).

51. Kent, W. J., Zweig, A. S., Barber, G., Hinrichs, A. S. & Karolchik, D. BigWig and BigBed: enabling browsing of large distributed datasets. Bioinformatics 26, 2204–2207 (2010).

52. Servant, N. et al. HiC-Pro: an optimized and flexible pipeline for Hi-C data processing. Genome Biol. 16, 259 (2015).

53. Bhattacharyya, S., Chandra, V., Vijayanand, P. & Ay, F. Identification of significant chromatin contacts from HiChIP data by FitHiChIP. Nat. Commun. 10, 4221 (2019).

54. Subramanian, A. et al. Gene set enrichment analysis: a knowledge-based approach for interpreting genome-wide expression profiles. Proc. Natl. Acad. Sci. U. S. A. 102, 15545– 15550 (2005).

55. Yardımcı, G. G. et al. Measuring the reproducibility and quality of Hi-C data. Genome Biol. 20, 1–19 (2019).

56. Harmston, N., Ing-Simmons, E., Perry, M., Barešić, A. & Lenhard, B. GenomicInteractions: An R/Bioconductor package for manipulating and investigating chromatin interaction data. BMC Genomics 16, 963 (2015).

57. Hagberg, A., Swart, P. & S Chult, D. Exploring network structure, dynamics, and function using networkx. https://www.osti.gov/biblio/960616 (2008).

58. Robinson, M. D. & Oshlack, A. A scaling normalization method for differential expression analysis of RNA-seq data. Genome Biol. 11, R25 (2010).

59. Buitinck, L., et al. API design for machine learning software: experiences from the scikit-learn project. arXiv [cs.LG] (2013).

60. Breiman, L. Mach. Learn. 45, 5–32 (2001).

61. Hunter. Matplotlib: A 2D Graphics Environment. 9, 90–95 (2007).

62. Virtanen, P., Gommers, R., Oliphant, T. E. & Haberland, M. Fundamental algorithms for scientific computing in python and SciPy 1.0 contributors. SciPy 1.0. Nat. Methods.

63. Liang, G. et al. Distinct localization of histone H3 acetylation and H3-K4 methylation to the transcription start sites in the human genome. Proc. Natl. Acad. Sci. U. S. A. 101, 7357– 7362 (2004).

64. Creyghton, M. P. et al. Histone H3K27ac separates active from poised enhancers and predicts developmental state. Proceedings of the National Academy of Sciences vol. 107 21931–21936 Preprint at 10.1073/pnas.1016071107 (2010).

65. Roadmap Epigenomics Consortium et al. Integrative analysis of 111 reference human epigenomes. Nature 518, 317–330 (2015).

66. Chen, Z. et al. Androgen Receptor-Activated Enhancers Simultaneously Regulate Oncogene TMPRSS2 and lncRNA PRCAT38 in Prostate Cancer. Cells 8, (2019).

67. Henriques, T., et al. Widespread transcriptional pausing and elongation control at enhancers. Genes & Development vol. 32 26–41 Preprint at 10.1101/gad.309351.117 (2018).

68. Breiman, L. Random Forests. Mach. Learn. 45, 5–32 (2001).

69. Lawrence, M. G., Lai, J. & Clements, J. A. Kallikreins on steroids: structure, function, and hormonal regulation of prostate-specific antigen and the extended kallikrein locus. Endocr. Rev. 31, 407–446 (2010).

70. Reményi, A., Schöler, H. R. & Wilmanns, M. Combinatorial control of gene expression. Nat. Struct. Mol. Biol. 11, 812–815 (2004).

71. Sproul, D., Gilbert, N. & Bickmore, W. A. The role of chromatin structure in regulating the expression of clustered genes. Nat. Rev. Genet. 6, 775–781 (2005).

72. Cosgrove, M. S., Boeke, J. D. & Wolberger, C. Regulated nucleosome mobility and the histone code. Nat. Struct. Mol. Biol. 11, 1037–1043 (2004).

73. Glont, S.-E., Chernukhin, I. & Carroll, J. S. Comprehensive Genomic Analysis Reveals that the Pioneering Function of FOXA1 Is Independent of Hormonal Signaling. Cell Rep. 26, 2558–2565.e3 (2019).

74. Le Dily, F. & Beato, M. Signaling by Steroid Hormones in the 3D Nuclear Space. Int. J. Mol. Sci. 19, (2018).

75. Karnuta, J. M. & Scacheri, P. C. Enhancers: bridging the gap between gene control and human disease. Hum. Mol. Genet. 27, R219–R227 (2018).

76. Fulco, C. P. et al. Activity-by-contact model of enhancer-promoter regulation from thousands of CRISPR perturbations. Nat. Genet. 51, 1664–1669 (2019).

77. Andersson, R. & Sandelin, A. Determinants of enhancer and promoter activities of regulatory elements. Nat. Rev. Genet. 21, 71–87 (2020).

78. Oudelaar, A. M. & Higgs, D. R. The relationship between genome structure and function. Nat. Rev. Genet. 22, 154–168 (2021).

79. Mumbach, M. R. et al. HiChIP: efficient and sensitive analysis of protein-directed genome architecture. Nat. Methods 13, 919–922 (2016).

80. Mumbach, M. R. et al. Enhancer connectome in primary human cells identifies target genes of disease-associated DNA elements. Nat. Genet. 49, 1602–1612 (2017).

81. Crispatzu, G. et al. The chromatin, topological and regulatory properties of pluripotency-associated poised enhancers are conserved in vivo. Nat. Commun. 12, 4344 (2021).

82. Cao, Z.-J. & Gao, G. Multi-omics single-cell data integration and regulatory inference with graph-linked embedding. Nat. Biotechnol. (2022) doi:10.1038/s41587-022-01284-4.

83. Pliner, H. A. et al. Cicero Predicts cis-Regulatory DNA Interactions from Single-Cell Chromatin Accessibility Data. Molecular Cell vol. 71 858–871.e8 Preprint at 10.1016/j.molcel.2018.06.044 (2018).

84. Thibodeau, A., Márquez, E. J., Shin, D.-G., Vera-Licona, P. & Ucar, D. Chromatin interaction networks revealed unique connectivity patterns of broad H3K4me3 domains and super enhancers in 3D chromatin. Sci. Rep. 7, 14466 (2017).

